# A simulated annealing algorithm for randomizing weighted networks

**DOI:** 10.1101/2024.02.23.581792

**Authors:** Filip Milisav, Vincent Bazinet, Richard F. Betzel, Bratislav Misic

## Abstract

Scientific discovery in connectomics relies on the use of network null models. To systematically evaluate the prominence of brain network features, empirical measures are compared against null statistics computed in randomized networks. Modern imaging and tracing technologies provide an increasingly rich repertoire of biologically meaningful edge weights. Despite the prevalence of weighted graph analysis in connectomics, randomization models that only preserve binary node degree remain most widely used. Here, to adapt network null models to weighted network inference, we propose a simulated annealing procedure for generating strength sequence-preserving randomized networks. This model outperforms other commonly used rewiring algorithms in preserving weighted degree (strength). We show that these results generalize to directed networks as well as a wide range of real-world networks, making them generically applicable in neuroscience and in other scientific disciplines. Furthermore, we introduce morphospace representation as a tool for the assessment of null network ensemble variability and feature preservation. Finally, we show how the choice of a network null model can yield fundamentally different inferences about established organizational features of the brain such as the rich-club phenomenon and lay out best practices for the use of rewiring algorithms in brain network inference. Collectively, this work provides a simple but powerful inferential method to meet the challenges of analyzing richly detailed next-generation connectomics datasets.

## INTRODUCTION

The connectome is a complex network that constitutes a comprehensive catalogue of the brain’s neural elements and their connections [102]. Numerous topological features of structural brain networks have been identified, including high clustering, short characteristic path length [9, 52, 57, 103, 122], and a rich-club of highly interconnected hub nodes [113, 115, 116]. These network features are consistently expressed across species [114], spatial scales, neuroimaging modalities, and tract-tracing technologies [101].

To quantify the unexpectedness of brain network features, network-based statistics are typically compared against null features computed in populations of randomized networks that preserve specific features of the empirical network [119]. Comparisons with randomized networks can then be used either for statistical inference or for benchmarking. In the case of statistical inference, a *p*-value for a particular network feature can be estimated by computing the proportion of randomized networks for which that feature has a more extreme magnitude than in the empirical network. In the case of benchmarking, a network feature can be normalized against the distribution of that feature in the population of randomized networks. For instance, the oft-studied small-world coefficient [53, 76, 122] and the rich-club coefficient [28, 82] are by definition normalized in such a manner. The most widely used network null model is degree-preserving rewiring (often referred to as Maslov-Sneppen rewiring; [70]), which uses edge swapping to disrupt the empirical network’s topology, but preserves ^*∗*^ its size (i.e., number of nodes), density (i.e., proportion of expressed edges), and binary degree sequence (i.e., number of edges incident to each node). Importantly, by selectively controlling for lower-order features, null models can be used to rule out the possibility that higher-order structures reflect random assemblies of simpler features [119].

However, with the advent of rich weighted networks— spanning up to six orders of weight magnitude [34, 35, 127]—and their increasing use over their simpler binary counterparts, there is a need for null models that accommodate weighted network statistics. For example, in diffusion-weighted MRI tractometry, edge weights are conventionally estimated using numbers of streamlines or fractional anisotropy [126]. More recently, numerous metrics have been developed to more directly quantify microstructural attributes of network edges [77], including neurite density [128], myelin [50, 68, 107, 108], axon diameter [3, 4], and axon cross-sectional area [30, 99]. Invasive methods in animal models also yield weighted networks, including tract tracing using fluorescent markers [66, 67, 69, 79], genetic labeling [24] and electron microscopy [51, 124]. Additionally, brain networks are increasingly reconstructed by means of comparing inter-regional similarity [10, 49], such as gene co-expression [14, 39], laminar profile covariance [84] or receptor similarity [48], again yielding biologically meaningful edge weight distributions. Therefore, next-generation connectomics requires new randomization algorithms that take edge weights into account.

Several network null models have been developed that preserve the weighted degree sequence (hereafter referred to as strength sequence) of empirical networks in addition to their binary degree sequence [41, 82, 88, 96, 104, 130]. Most of these models are sampling methods based on maximum-likelihood estimation of a network probability distribution [41, 96, 104]. Here, we do not consider these models for two reasons. First, they only satisfy the degree sequence constraints on average across the complete network ensemble, but not for each individual sampled network [26, 89]. Second, these models do not maintain the empirical network’s weight distribution [89]. Furthermore, for these reasons, they are seldom practically used in network neuroscience.

### Algorithm 1

Strength sequence-preserving network randomization

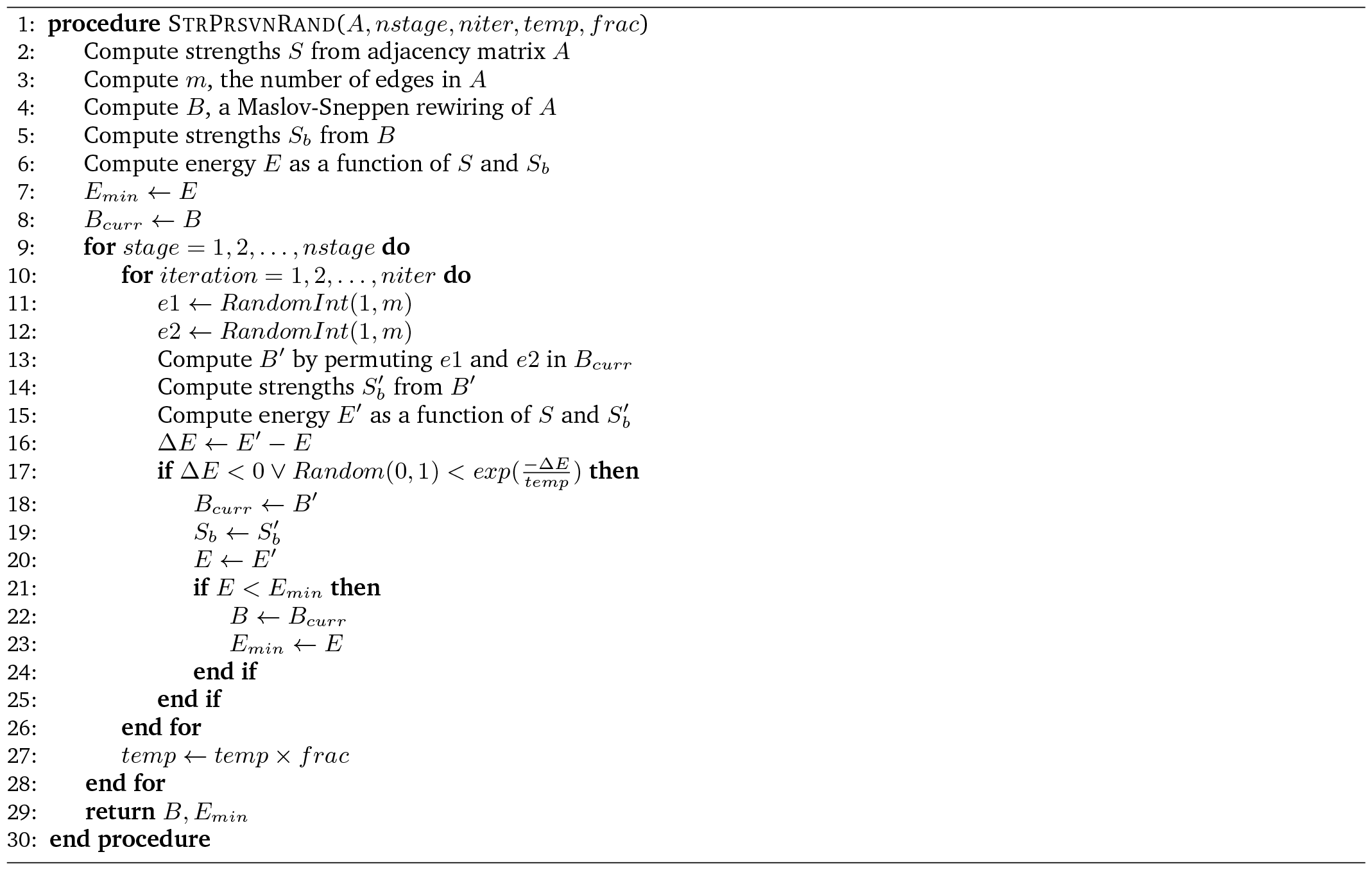

Here we present an algorithm that addresses these limitations by preserving an empirical network’s weight distribution, degree sequence, and strength sequence for each randomized network instance. Contrary to other strength-preserving models, this procedure does not require any analytical derivations, instead building on classic rewiring techniques already commonplace in the field of network neuroscience and network science more generally. This randomization technique reconfigures weight placement atop the binary scaffolding of a rewired network to match the empirical network’s strength sequence using simulated annealing, a probabilistic algorithm that approximates the global minimum of a given function [59, 60]. Simulated annealing is a powerful and versatile optimization technique with wide-ranging applications. Moreover, it is particularly advantageous when dealing with large combinatorial search spaces, making it a prime candidate for solving network modeling problems [1, 19, 23, 26, 57, 74, 83, 86, 89, 100, 120].

In this report, we benchmark the performance of the simulated annealing procedure against another rewiring algorithm for strength sequence-preserving randomization (hereafter referred to as the Rubinov-Sporns algorithm; [92]), as well as the classic Maslov-Sneppen degree-preserving rewiring model [70]. In parallel, we introduce novel tools for assessing null network variability, a seldom considered [8, 61, 73], but important evaluation step when comparing network null models.

## RESULTS

The results are organized as follows. We first introduce a simple and versatile rewiring algorithm based on simulated annealing that preserves the strength sequence and weight distribution of the empirical network. We then systematically evaluate the capacity of the algorithm to preserve the strength sequence compared to two existing state-of-the-art procedures conventionally used in the field. We also introduce morphospace representation as a graphical tool for the assessment of variability and feature preservation in null networks. Finally, we consider the weighted rich-club phenomenon as a practical example of the influence a network null model can exert on weighted network inference. Throughout the analyses, we extensively test algorithm performance across multiple independent datasets and a variety of preprocessing choices to capture null model behavior in different contexts.

Briefly, we consider three rewiring algorithms (Fig. 1; for a detailed description, see *Methods*). In the classic Maslov-Sneppen algorithm, pairs of edges are randomly swapped, and weights are “carried” with their respective edges [70] (Fig. 1a). The Rubinov-Sporns algorithm, building on the output of the Maslov-Sneppen algorithm, attempts to preserve the strength sequence by sampling each edge weight, in pseudorandom order, from the original edge weight distribution using a rank-matching procedure [92] (Fig. 1b). Finally, in the simulated annealing procedure, also building on the output of the Maslov-Sneppen algorithm, randomly selected pairs of edge weights are permuted either if they lower the energy of the system (mean squared error between the strength sequences of the empirical and the randomized networks) or if they meet a probabilistic acceptance criterion. This allows permutations which can increase the energy of the system but prevents it from getting stuck in a local minimum (Fig. 1c). Importantly, the procedure is also applied on the Maslov-Sneppen rewired network. Full details of the algorithm are shown in Algorithm 1 and a Python implementation is openly available as part of the netneu-rotools package (https://netneurotools.readthedocs.io/en/latest/api.html#module-netneurotools.networks).

**Figure 1.**
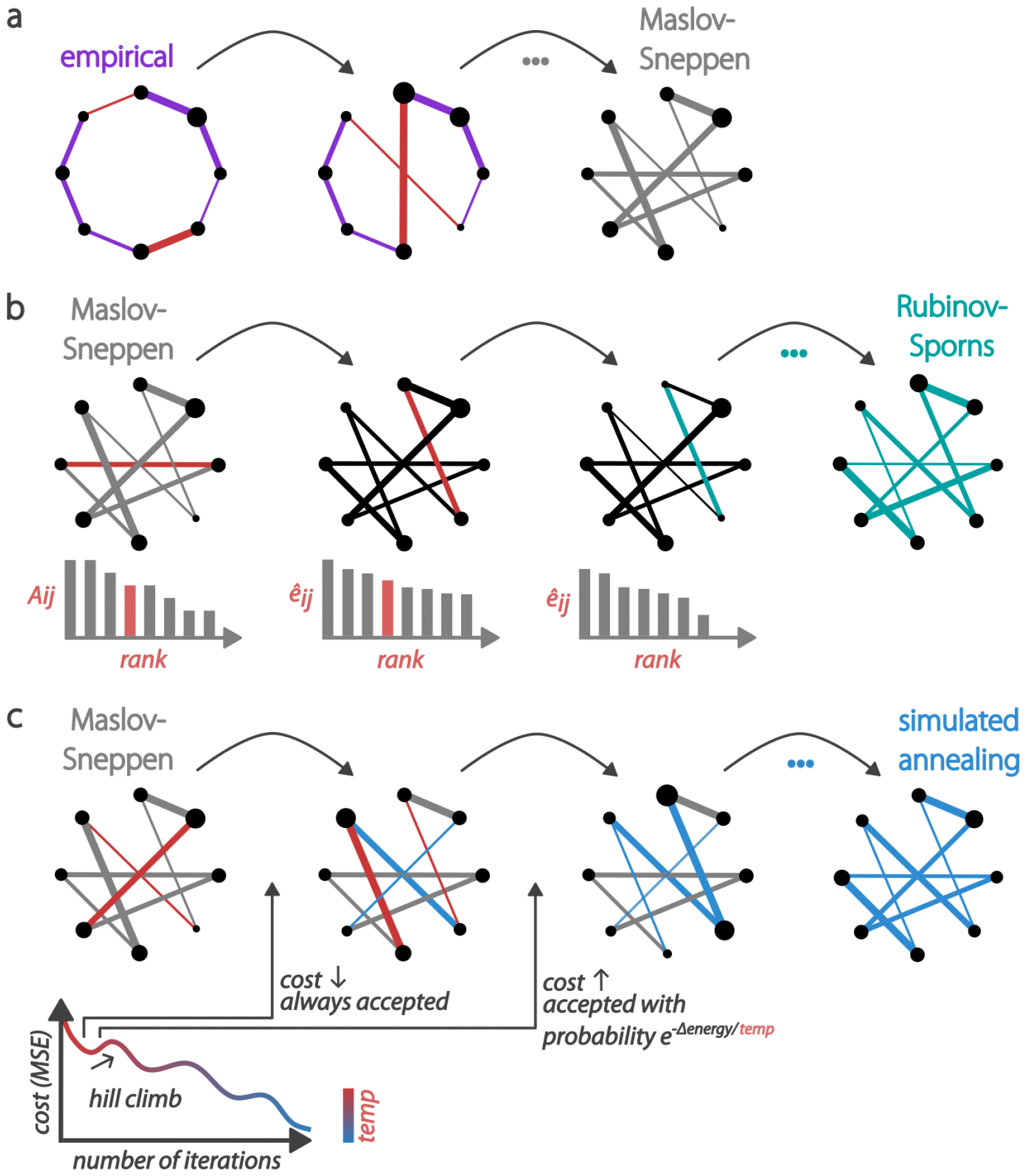
Rewiring algorithms for generating strength sequence-preserving randomized null networks. (a) Maslov-Sneppen degree-preserving rewiring [70]: pairs of edges (red) are randomly swapped, disrupting the network’s topology, but preserving its size, density, and degree sequence. Edge width represents weight and node size represents strength. (b) Rubinov-Sporns strength sequence-preserving randomization algorithm [92]: using the Maslov-Sneppen rewired network, (1) the randomized network is instantiated with zeros (*Â*_*ij*_ = 0; in black); (2) the original edge weights (*A*_*ij*_) are ranked by magnitude (left); (3) the edges of the randomized network are ranked by their expected magnitude (*ê*_*ij*_ ; middle); (4) a random edge is selected in the randomized network and its weight is set to the original edge weight of the same rank (both edges are depicted in red); (5) edges are re-ranked and the procedure is repeated, resulting in the Maslov-Sneppen rewired network, with edge weights permuted to approximate the empirical network’s strength sequence (right). *ê*_*ij*_ *∝* (*s*_*i*_ *−* Σ_*u*_*Â*_*iu*_)(*s*_*j*_ *−* Σ_*u*_*Â*_*ju*_), where *s*_*i*_ represents the strength of node *i* in the empirical network and Σ_*u*_*Â*_*iu*_ is the sum of the weights of assigned edges incident to node *i* in the randomized network. In the middle networks, edge width represents expected weight magnitude (black or red) or assigned weight (teal) and node size represents residual strength (*s*_*i*_ *−* Σ_*u*_*Â*_*iu*_). (c) Strength sequence-preserving randomization via simulated annealing: using the Maslov-Sneppen rewired network, randomly selected pairs of edge weights (red) are permuted either if they lower the energy of the system (mean squared error between the strength sequences of the empirical and the randomized networks) or if they meet the probabilistic Metropolis acceptance criterion, depending on the temperature of the system.

Experiments are performed in two publicly available diffusion-weighted MRI datasets, acquired using different protocols (diffusion spectrum imaging; DSI, and high angular resolution diffusion imaging; HARDI), parcellations (anatomical and functional) and parcellation resolutions (low and high in each dataset). The first sample (LAU) consists of DSI data acquired in *n* = 70 participants (source: Lausanne University Hospital [46]; see *Methods* for detailed procedures). The second sample consists of HARDI data acquired in *n* = 327 participants (source: Human Connectome Project – HCP [117]; see *Methods* for detailed procedures). For both datasets, group-representative weighted structural networks were built using a distance-dependent consensus-based thresholding procedure [13, 75], resulting in a total of 4 empirical group-consensus networks (LAU - low res, LAU - high res, HCP - low res, HCP - high res) on which the main analyses were conducted (but see section “Strength-preserving randomization in individual networks” for analysis of individual participant connectomes). Here, results are visualized in the high resolution HCP dataset but exhaustive figures are provided as *Supplementary Information*.

### Benchmarking strength sequence preservation

We benchmark the performance of the randomization algorithms by generating 10 000 null networks for each empirical brain network. To characterize the performance of the null models in preserving strength sequence—the ordered set of strengths in which each node is associated to a specific strength—we plot empirical network strengths against strengths of the randomized networks for all 10 000 nulls (Fig. 2a, S4). We also compute Spearman rank-order correlation coefficients between empirical and “randomized” strengths. Across all four empirical brain networks, the simulated annealing algorithm yields near perfect fits (LAU - low res: *M ≈* 0.999, *SD ≈* 0.001, LAU - high res: *M ≈* 0.996, *SD ≈* 0.002, HCP - low res: *M ≈* 1.0, *SD ≈* 3.75 *×* 10^*−*7^, HCP - high res: *M ≈* 1.0, *SD ≈* 1.88 *×* 10^*−*7^). It also results in larger correlation coefficients than the Rubinov-Sporns algorithm (*p ≈* 0, effect size = 100% for all empirical networks, two-tailed, Wilcoxon–Mann–Whitney two-sample rank-sum test), which itself results in larger coefficients than the Maslov-Sneppen algorithm (*p ≈* 0, effect size = 100% for all empirical networks, two-tailed, Wilcoxon–Mann–Whitney two-sample rank-sum test). This shows that the simulated annealing algorithm generates randomized networks with the most veridical strength sequences. In the *Supplementary Information*, we further investigate null model calibration (section “Null model calibration”) and provide sensitivity analyses examining the effect of an alternative objective function (section “Alternative objective function”) and weight log-transformation on simulated annealing performance (section “Log-transformation”).

**Figure 2.**
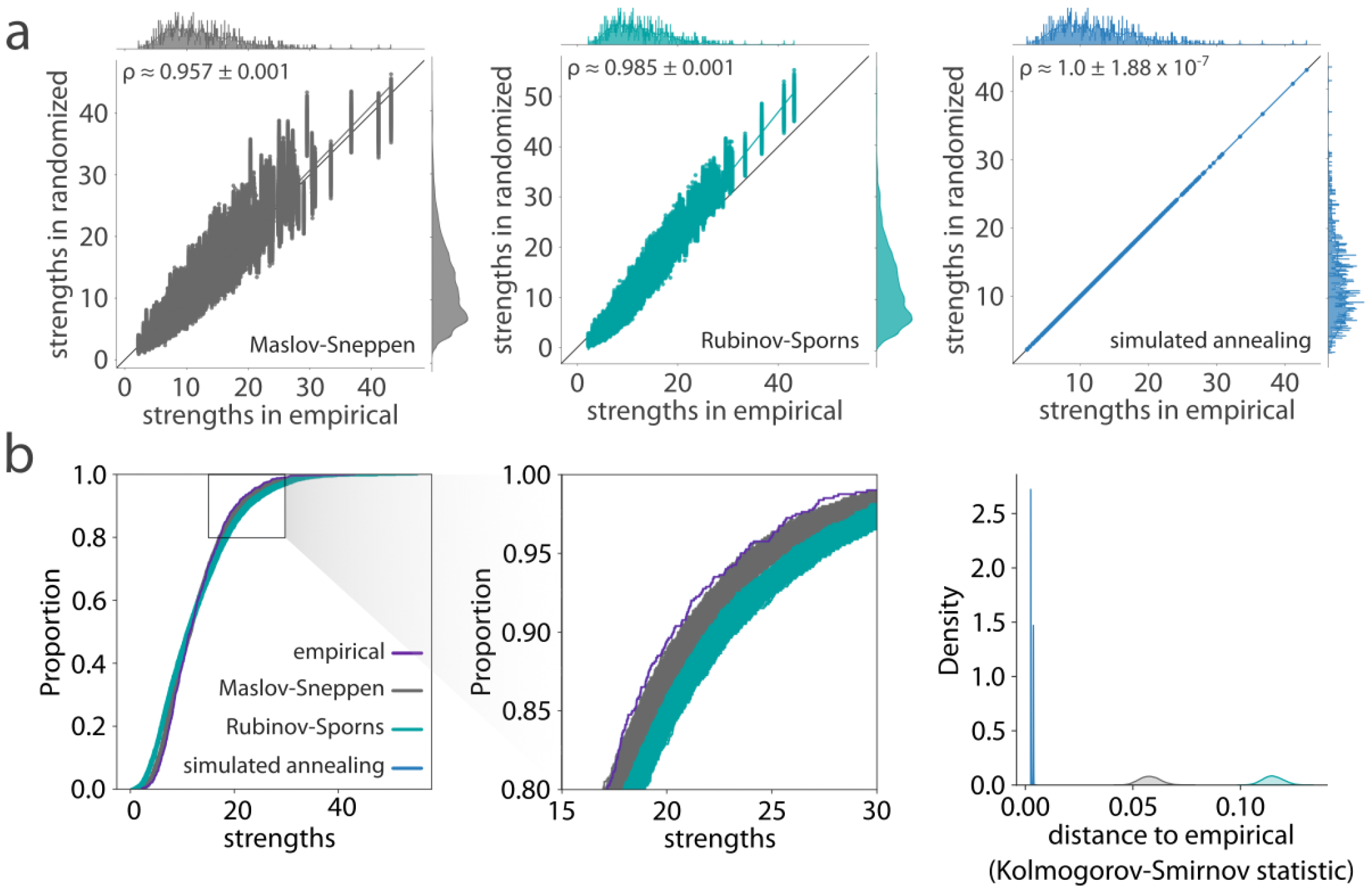
Benchmarking strength preservation. (a) Scatter plots of strengths of the empirical (abscissa) and randomized (ordinate) networks for all 10 000 null networks, where each point represents a brain region. Marginal distribution histograms are shown on the top and right axes. Mean and standard deviation across 10 000 Spearman rank-order correlation coefficients are provided as insets. Data points and histograms appear in grey for the Maslov-Sneppen algorithm, teal for the Rubinov-Sporns algorithm, and blue for the simulated annealing algorithm. Linear regression lines (colored) were added for visualization purposes. The identity line (black) is provided as reference. (b) Strength cumulative distribution functions (left) and density plots representing Kolmogorov-Smirnov statistics obtained by comparing the strength distribution of the empirical network with that of the randomized networks (right). Cumulative and probability density function curves are shown in grey for the Maslov-Sneppen algorithm, teal for the Rubinov-Sporns algorithm, and blue for the simulated annealing algorithm. The original cumulative distribution function is depicted in indigo and almost perfectly overlays all 10 000 cumulative distribution functions obtained via simulated annealing, effectively hiding them.

### Benchmarking strength distribution preservation

While we have established, using rank-based methods, that the simulated annealing algorithm outperforms other randomization techniques in preserving the empirical network’s strength sequence, we have not quantified how well the different models preserve the strength distribution. The level to which the empirical strength distribution is preserved in a null network is crucial, because it ensures an accurate representation of influential graph features, such as hubs, whose significance is intricately tied to characteristics of the distribution.

To assess the goodness of fit between the strength distributions of the empirical and the randomized structural networks, we superimpose their cumulative distribution functions (Fig. 2b; left, Fig. S5; top). Across all datasets, the curves produced via simulated annealing show the best match to the empirical strength cumulative distribution function with almost perfect superposition. Furthermore, the curves obtained using the Rubinov-Sporns and the Maslov-Sneppen algorithms show considerably more variability across null networks as shown by their wider spread, recapitulating previously observed patterns of underestimation and overestimation across datasets (see section “Null model calibration”). To confirm these observations quantitatively, we compute Kolmogorov-Smirnov (KS) test statistics between the cumulative strength distributions of the empirical and each randomized network, measuring the maximum distance between them (Fig. 2b; right, Fig. S5; bottom). Across all datasets, the simulated annealing algorithm outperforms the other two null models with significantly lower KS statistics (*p ≈* 0, effect size = 100% for all two-tailed, Wilcoxon–Mann–Whitney two-sample rank-sum tests). Furthermore, in the HCP dataset and the higher resolution Lausanne network, the Rubinov-Sporns algorithm generated cumulative strength distributions with slightly worse correspondence to the empirical distribution than the cumulative strength distributions yielded by the Maslov-Sneppen algorithm (LAU - high res: *p <* 10^*−*176^, effect size = 61.58%, HCP: *p ≈* 0, effect size = 100% for all empirical networks, two-tailed, Wilcoxon–Mann–Whitney two-sample rank-sum test).

As an illustration, we consider whether the nulls generated by different algorithms recapitulate fundamental characteristics associated with the empirical strength distribution. Namely, we focus on the heavy-tailedness of the strength distribution (i.e., does the null network also have a heavy-tailed strength distribution, suggesting the presence of hubs?) and the spatial location of high-strength hub nodes. We assess heavy-tailedness and identify hubs using the nonparametric procedure outlined in [40, 56] (see *Methods* for more details). Briefly, this procedure entails identifying hubs as the right tail outliers of the strength distribution. The proportion of outliers, i.e., right-tailedness, is then compared to the parameter-invariant right-tailedness of the exponential distribution. Heavy-tailedness is detected if the empirical right-tailedness exceeds the exponential right-tailedness, that is, if the tail decay of the empirical distribution is subexponential [38, 56].

Considering the high-resolution Lausanne dataset as an example, we find that the empirical group-consensus network has a heavy-tailed strength distribution, with 2% of the nodes identified as hubs (Fig. S6b); Fig. S6a shows their spatial location in red. By comparison, across 10 000 realizations, the Maslov-Sneppen algorithm only recapitulates heavy-tailedness 0.03% of the time and the spatial locations of hubs do not recapitulate the empirical map. The Rubinov-Sporns algorithm does better and detects heavy-tailedness in all realizations, but identifies only 1.4% of nodes as hubs on average (compared to 2% in the empirical network). Finally, the simulated annealing algorithm detects heavy-tailedness in all realizations and identifies 1.96% of nodes as hubs on average, providing the closest match to the empirical network. We also assess how well each algorithm recovers the correct spatial location of these hubs using z-scored Rand indices between the empirical and the null hub assignments and find that simulated annealing also outperforms the other algorithms in recapitulating hub identity (Fig. S6c; *p ≈* 0, effect size *≈* 100% for both two-tailed, Wilcoxon–Mann–Whitney two-sample rank-sum tests).

### Null network ensemble variability

Generating rewired null networks for the purpose of null hypothesis testing or normalization is ubiquitous in network neuroscience, and in network science more generally [119]. Yet the features derived from ensembles of rewired nulls are typically averaged or summarized, without considering the variability across network realizations. Whether all rewiring algorithms produce ensembles of null networks that vary to the same extent is unknown. Do different algorithms yield null networks with comparable architectural features? Do different algorithms sample the space of possible null networks differently? To assess variability of network surrogate realizations, we embedded the 10 000 nulls generated for each randomization algorithm and each of the four empirical networks in a two-dimensional morphospace spanned by two global network statistics: characteristic path length and clustering (Fig. 3a) [5, 6, 29, 44, 45]. Note that this method could have been applied with any other network statistic; we chose characteristic path length and clustering because of their wide use in the network science literature, notably as a means to study small-worldness [9, 52, 57, 103, 122].

**Figure 3.**
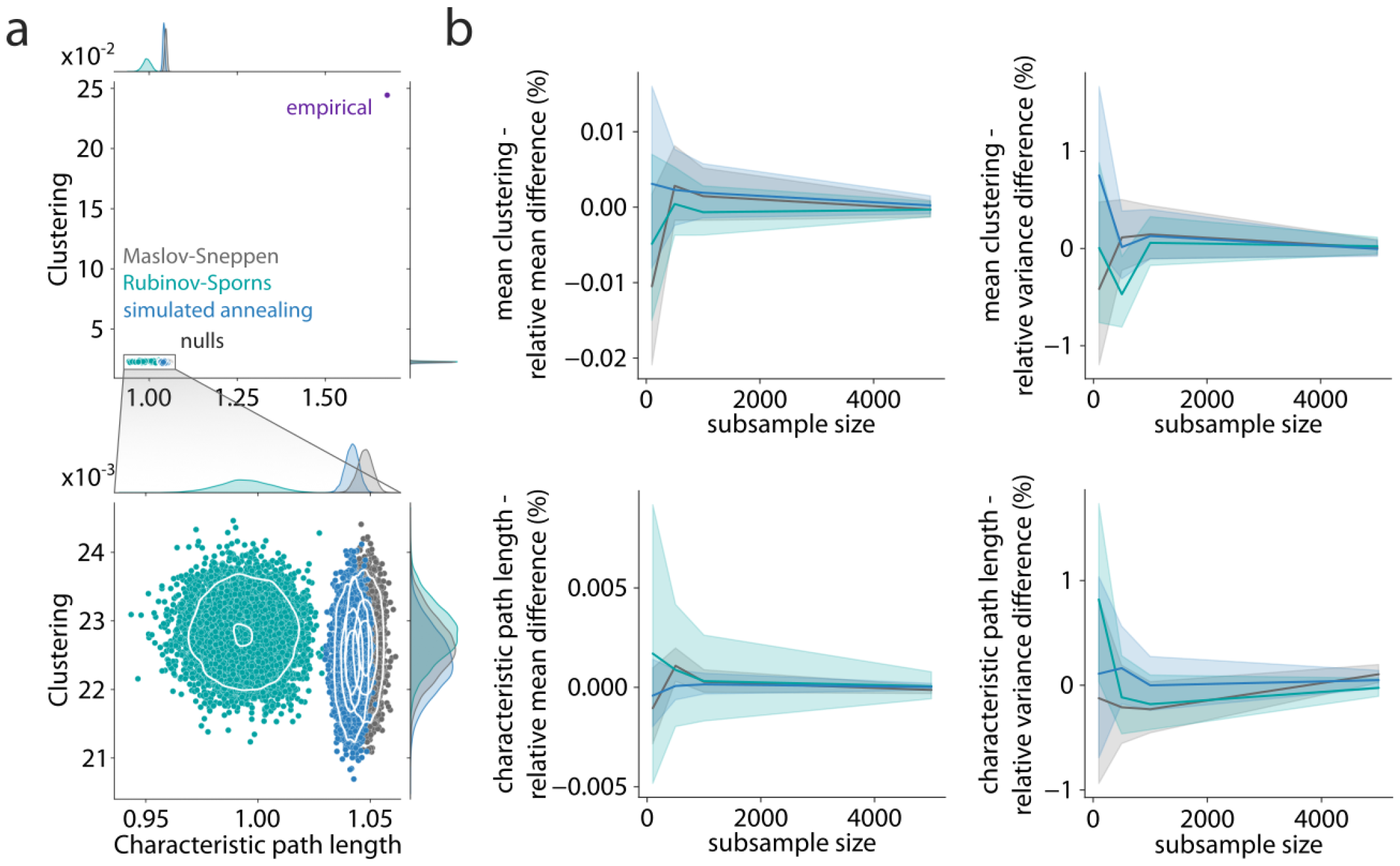
Morphospace of null network ensembles. (a) Morphospaces spanned by characteristic path length and clustering. Marginal distribution histograms are shown on the top and right axes. Data points corresponding to randomized null networks generated by the simulated annealing algorithm appear in blue; those resulting from the Rubinov-Sporns algorithm appear in teal; and Maslov-Sneppen rewired networks are shown in grey. The empirical group-consensus structural network is depicted in indigo. The bottom panel consists in a zoomed-in view of the clusters of randomized networks appearing in the top panel. Contour levels are drawn using a Gaussian kernel density estimate and delineate iso-proportions of the density. (b) Trajectories of relative difference in mean clustering (top left), clustering variance (top right), mean characteristic path length (bottom left) and characteristic path length variance (bottom right) between the full null population (*N* = 10 000) and subsamples of increasing size (*n* ∈ {100, 500, 1000, 5000}). Colored lines and shaded bands represent mean and 95% bootstrapped confidence interval (1000 samples).

Across datasets, we observe that randomized networks generally occupy the same portion of the morphospace relative to the empirical networks (Fig. 3a, top). This indicates that despite the further constraints in edge weight placement imposed upon the strength sequence-preserving algorithms, they produce similar patterns to the classic Maslov-Sneppen degree-preserving rewiring. While this is true when comparing the position of randomized networks to that of the empirical network, zooming-in on the region of the morphospace occupied by the null networks (Fig. 3a, bottom) reveals important differences in their variability.

Null networks derived using simulated annealing organized in patterns similar to those of the Maslov-Sneppen rewired networks. Namely, the simulated annealing ensemble retains a similar shape, similar density distribution (as shown by the contour lines) and often occupies a position close to the Maslov-Sneppen ensemble that remains consistent across datasets (Fig. 3a, S7). By comparison, the Rubinov-Sporns ensemble is less consistent across datasets and in the HCP dataset, yields large density distributions in which many realizations are disproportionately concentrated in the middle. Interestingly, simulated annealing ensembles generally show similar clustering but slightly lower characteristic path length than their strictly degree-preserving counterpart (*p ≈* 0 for all empirical networks, two-tailed, Wilcoxon–Mann–Whitney two-sample rank-sum test, LAU - low res: effect size = 97.66%, LAU - high res: effect size = 100%, HCP - low res: effect size = 89.11%, HCP - high res: effect size = 90.26%). While this shows that the simulated annealing algorithm yields networks that are more dissimilar to the empirical network in terms of their characteristic path length as compared to Malsov & Sneppen rewiring, it also identifies them as slightly more stringent benchmarks when assessing how unexpectedly low an empirical network’s characteristic path length is, such as when computing the small-world coefficient [53, 122].

Another question that naturally emerges when using network null models is how many nulls to generate. While the scaling behavior of a network feature’s null distribution probably varies depending on the feature at hand, morphospaces might provide insight into the question by summarizing global aspects of a network’s architecture. In Fig. S8, we consider a subsample of 100 nulls out of the 10 000 generated. We see that the same patterns of null network ensemble organization already seem to emerge with only 100 nulls, sampling a similar extent of the morphospace. To quantify the scaling behavior of the morphospace, we consider a range of increasing sub-sample sizes (*n ∈* 100, 500, 1000, 5000). For each randomization algorithm and sub-sample size, we draw 1000 random samples and compute the relative difference between each sample’s global network statistics (mean and variance across nulls of mean clustering and characteristic path length) and the statistics obtained in the full ensemble of 10 000 nulls. In Fig. 3b and S9, we show the scaling behavior of the morphospace’s global network statistics as a function of null sample size. Interestingly, we find that even at the lowest sample size, relative differences do not exceed 1% for any of the statistics. Furthermore, relative differences rapidly converge to even lower values with increasing sample size. Therefore, we find that only a small number of nulls is necessary to adequately approximate the null distribution of these global network features. Altogether, these results show that explicitly considering how the space of possible null realizations is sampled is important yet often overlooked. Additionally, in the *Supplementary Information*, we analyze morphospace trajectories, relating energy and morphospace position throughout the simulated annealing procedure (section “Morphospace trajectories”).

### The weighted rich-club phenomenon

To illustrate how the choice of a network null model can have important ramifications for network inference, we consider the weighted rich-club phenomenon in connectomics. A rich-club is characterized as a group of high-degree nodes (rich nodes) that exhibit a greater number of interconnections than would be anticipated by chance [28, 129]. In this study, we go beyond the conventional definition of rich-club and incorporate a weighted measure to assess the relative strength of connections among rich nodes, referred to as the weighted rich-club coefficient [82]. This measure is evaluated at various threshold degree values, used to define the rich nodes (see *Methods* for more details). Considering that high-degree nodes are more likely to be interconnected, the (weighted) rich-club coefficient is commonly normalized against a null rich-club coefficient averaged across an ensemble of randomized null networks. Although the conventional Maslov-Sneppen degree-preserving rewiring method is frequently employed in generating the null network population, it does not factor in the effect of the weighted degree sequence in the computation of the weighted rich-club coefficient. Here, to contrast inferences obtained using different null models, we compute the weighted rich-club coefficient in each of the 10 000 null networks generated for each model under study. We then compute the normalized weighted rich-club coefficient using the average coefficient across nulls for each model and assess significance by deriving a p-value as the proportion of null coefficients that are greater than the empirical coefficient.

Interestingly, we find that, for all empirical networks, using simulated annealing-derived null networks yields larger normalized rich-club ratios than using Rubinov-Sporns or Maslov-Sneppen randomization (Fig. 4, left, S11, top; *p <* 0.01 for all two-tailed, Wilcoxon–Mann–Whitney two-sample rank-sum tests).

**Figure 4.**
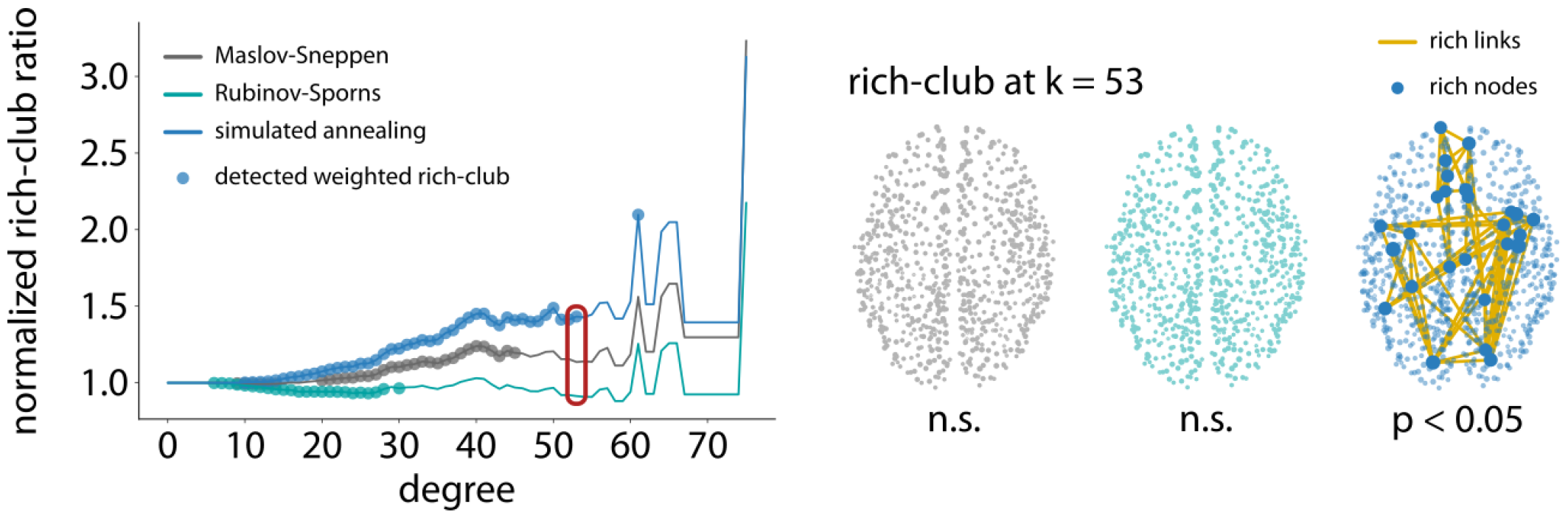
The weighted rich-club phenomenon. (a) Left: Normalized rich-club ratio as a function of the degree threshold used to define rich nodes. Lines are colored by the null algorithm used (Maslov-Sneppen in grey, Rubinov-Sporns in teal, and simulated annealing in blue). Colored points indicate significance at the Bonferroni-corrected threshold of *p <* 0.05, indicating that a weighted rich-club was detected. Right: Example empirical weighted rich-club detected at *k* = 53, with *k* corresponding to degree. Node size is proportional to strength. Only the simulated annealing algorithm detects a significant weighted rich-club at *k* = 53. Rich nodes are enlarged to showcase their spatial location.

Importantly, using the Maslov-Sneppen algorithm, no weighted rich-club is identified in the Lausanne dataset and only the simulated annealing algorithm identifies a weighted rich-club in the low-resolution version. This result might seem counter-intuitive given that null networks that embody more aspects of the empirical network are generally seen as more conservative. However, this is not a general rule and a specific result needs to be interpreted against the backdrop of the specific analysis and null constraints at hand. Here, we posit that the differences in normalized rich-club coefficient observed between models is due to an overestimation of strength in high-degree nodes. We further suggest that this difference between empirical strength and strength expected based on degree might be due to high-degree (rich-club) nodes being interconnected by a preponderance of low-weight long-range connections [115]. In the *Supplementary Information*, we verify these hypotheses, relating the choice of network null model to weighted rich-club inference and the spatial embedding of rich connections (section “Weighted rich-club inference and geometry”). Altogether, these results show how contrasting nulls embodying hierarchical constraints can provide layered in-sights into brain network organization. More broadly, the choice of network null model can yield fundamentally different inferences about hitherto established phenomena.

### Computational cost

The results so far show that simulated annealing out-performs alternative network null models. Given the pervasive assumption that simulated annealing is time consuming, we next sought to benchmark the computational cost associated with each procedure.

Unlike the other two algorithms, simulated annealing naturally involves a tradeoff between computational cost and performance. Namely, the user specifies the number of iterations per annealing stage—more iterations result in better solutions but this comes at the cost of added execution time. Fig. S12a (left) shows that MSE is logarithmically reduced with the number of iterations per annealing stage (blue), with a concomitant linear increase in execution time (yellow). To illustrate the benefit of added iterations, Fig. S12a (right) shows strength sequence fits at a small (1000) and large (100 000) number of iterations. Importantly, simulated annealing’s flexible nature allows it to reach arbitrarily optimal solutions with a sufficiently slow cooling schedule [60, 78].

Given that simulated annealing is more computationally intensive, how feasibly can it be applied to more fine-grained networks with a greater number of edges? Would users be forced to use the Maslov-Sneppen or Rubinov-Sporns algorithms in those instances? Fig. S12b shows execution time for the three null model algorithms run on empirical networks with increasing density. First, we observe that the Maslov-Sneppen and Rubinov-Sporns algorithms have considerably shorter execution time. However, we find that simulated annealing scales well, showing a near-flat relation with increasing density. By comparison, the Rubinov-Sporns algorithm shows a slightly worse scaling behavior, characterized by a steeper slope. Finally, the execution time of Maslov-Sneppen rewiring is the most sensitive to network density. Overall, these results show the flexibility of the simulated annealing procedure: despite greater computational cost, simulated annealing can be tuned to achieve even better performance, and it scales well to more detailed networks.

### Strength-preserving randomization in individual networks

So far we have only used empirical networks derived from a collation of individual-specific data. Therefore, how the use of the randomization algorithms under study translates to participant-level designs remains unknown. Notably, how does empirical variability compare to null variability and how much inter-individual variability is preserved in null networks?

To address these questions, we consider 69 density-matched individual structural connectivity networks from the low-resolution Lausanne dataset. We then generate 100 null networks per algorithm and empirical network and benchmark strength-preservation performance across randomization algorithms, but now in each individual network. We use an energy threshold for the simulated annealing algorithm to ensure a similar quality of strength reconstruction across networks (see *Methods* for more details). Fig. S13a shows the distributions of Spearman correlation coefficient between strength sequences in empirical and randomized networks. Again, we find that the simulated annealing algorithm outperforms other null models in preserving strength sequence (*p ≈* 0 for both two-tailed, Wilcoxon–Mann–Whitney two-sample rank-sum tests, simulated annealing - Rubinov-Sporns: effect size = 92.57%, simulated annealing - Maslov-Sneppen: effect size = 100%).

Next, to contrast empirical and null variability, we embed both empirical and null networks in the same morphospace (Fig. S13b). We observe that the empirical networks span a larger portion of the morphospace than the null networks, especially in terms of clustering. This might seem counter-intuitive given the pervasive idea that null brain networks are generically “random” and that as such, they should account for a larger space of possible realizations than empirical brain networks. However, random networks constitute a class of networks in their own right with their own characteristic features, such as low clustering and short characteristic path length [2, 122], and therefore we should not expect them to show more variability than empirical networks when assessed on that basis. Furthermore, random networks are also defined based on specific structural constraints (size, density, weight distribution, degree/strength sequence in this case).

To further explore the question of inter-individual variability, we focus on the null network ensembles. In Fig. S13, we color nulls by the algorithm used to generate them (Fig. S13d) and the empirical networks from which they are built (Fig. S13e). We observe that despite a considerably reduced variability, participant identity seems to dominate the organization of null networks, mimicking the empirical pattern of inter-individual variability. To confirm this intuition, we measure inter-individual Euclidean distance between participants in the morphospace for the empirical and null networks. We then compare algorithm-wise distributions of Spearman correlation coefficients between empirical and null Euclidean distances (Fig. S13c). We find that simulated annealing provides a better preservation of inter-individual morphospace distances than both the Rubinov-Sporns (*p <* 10^*−*9^, effect size = 75.23%, two-tailed, Wilcoxon–Mann–Whitney two-sample ranksum tests) and Maslov-Sneppen (*p <* 10^*−*10^, effect size = 77.03%, two-tailed, Wilcoxon–Mann–Whitney two-sample rank-sum tests) algorithms. This indicates that strength-preserving nulls also preserve inter-individual patterns of global weighted network features.

### Strength-preserving randomization for directed networks

An advantage of randomizing networks using simulated annealing is that the optimization algorithm can be readily applied to preserve multiple network features. A straightforward extension is the preservation of in- and out-strength sequences in directed networks [89]. This is important because axonal projections are fundamentally directed, and numerous techniques can be used to reconstruct afferent and efferent connections, such as tract tracing and genetic labeling [22, 58, 64, 65]. Here we simply reformulate the objective function to account for the reconstruction error associated separately with instrength and out-strength, allowing us to effectively recapitulate in- and out-strength sequences.

Fig. 5 (top) shows the directed, weighted wiring diagrams of the drosophila (fruit fly) [24, 98, 125], mouse [79, 89], rat [17], and macaque [95] along with scatter plots of strengths in the empirical and the randomized networks, separately for in- and out-strengths (bottom). Similarly to the results observed in the Lausanne dataset (see Fig. S4 and sections “Null model calibration” and “Log-transformation”), we observe some low-strength inaccuracies in the drosophila and mouse connectomes. However, they only have a minimal effect on strength reconstruction, as shown by the consistently high correlation coefficients obtained in all networks.

**Figure 5.**
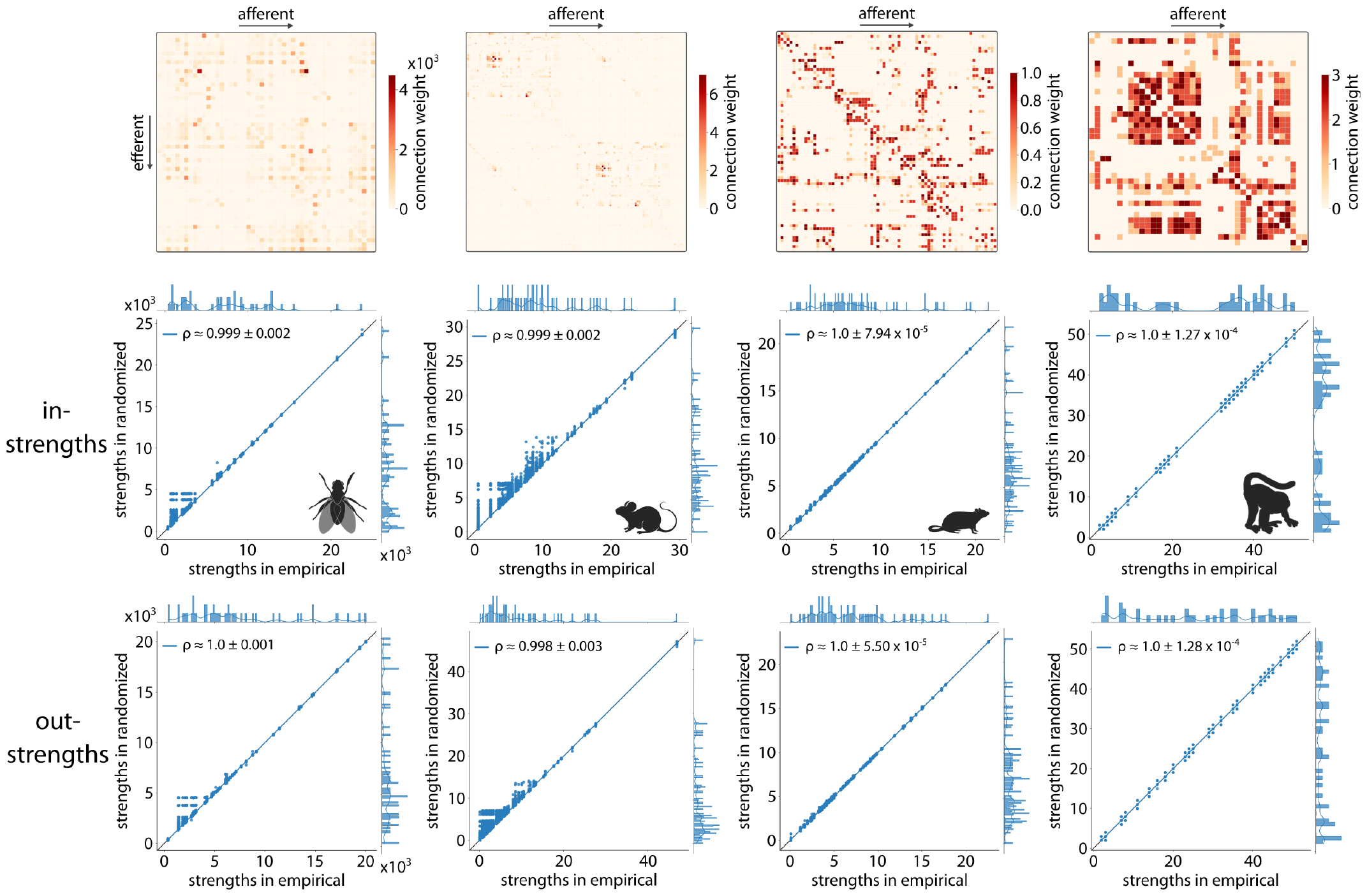
Strength-preserving randomization for directed networks. Top: Wiring diagrams for the drosophila, mouse, rat, and macaque connectomes (from left to right). Bottom: Scatter plots of strengths of the empirical (abscissa) and simulated annealing-derived networks (ordinate) for all 10 000 nulls, where each point represents a brain region. In-strengths and out-strengths are considered separately. Marginal distribution histograms are shown on the top and right axes. Mean and standard deviation across 10 000 Spearman rank-order correlation coefficients are provided as insets. Linear regression lines (blue) were added for visualization purposes. The identity line (black) is provided as reference.

### Strength-preserving randomization in real-world networks

Until now, we have only considered networks that represent inter-regional connectivity in brains. Yet the simulated annealing algorithm presented here is generic and can be readily applied to other classes of complex networks. For completeness, we benchmark strength-preservation performance in a dataset of 37 real-world weighted networks [42, 80]. The networks span multiple domains, with 28 networks representing social data, 1 network representing informational data, 2 networks representing biological data, 1 network representing economic data, and 5 networks representing transportation data. The networks are also diverse in terms of their basic features: network sizes range from 13 to 1707 nodes and densities range from 0.3% to 78.31% of connections present.

Fig. 6a shows the networks embedded in a low-dimensional morphospace, illustrating the breadth of network morphologies in the dataset. Fig. 6b shows the distributions of Spearman correlation coefficient between strength sequences in empirical and randomized networks. As with brain networks, simulated annealing consistently outperforms both the Rubinov-Sporns (effect size = 83.23%) and Maslov-Sneppen (effect size = 94.82%) algorithms (*p ≈* 0 for both two-tailed, Wilcoxon–Mann–Whitney two-sample rank-sum tests), with a peak near *ρ* = 1 for most networks in the dataset. This result demonstrates the broad utility of the approach beyond neuroscience.

**Figure 6.**
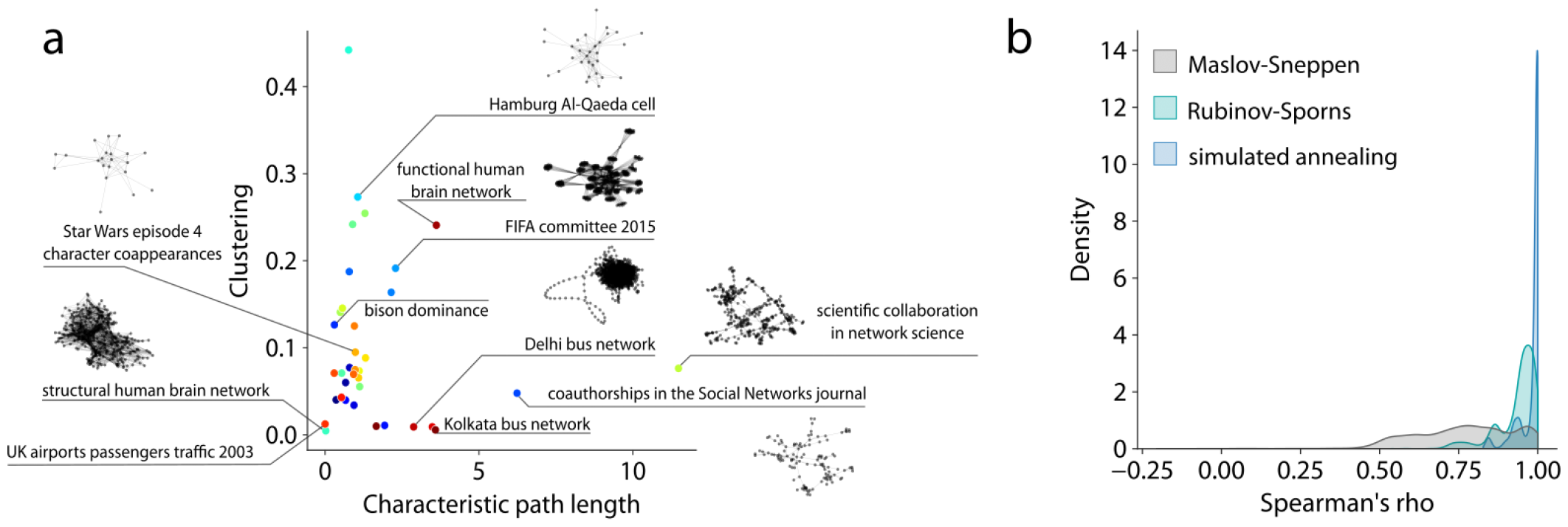
Strength-preserving randomization in real-world networks. (a) Morphospace spanned by characteristic path length and clustering in which all the weighted real-world networks under study are embedded. A subset of networks are identified and visualized using spring-embedding to showcase domain and morphological variability. (b) Density plot representing Spearman correlation coefficients between strength sequences in empirical and randomized networks derived using the Maslov-Sneppen algorithm (grey), Rubinov-Sporns algorithm (teal), and simulated annealing algorithm (blue).

## DISCUSSION

In the current report, we formally characterize a simulated annealing procedure for generating strength sequence-preserving randomized null networks. We benchmark this method against another previously used algorithm for strength-preserving weight reconfiguration [92], as well as the classic Maslov-Sneppen degree-preserving rewiring [70]. We find that the simulated annealing algorithm exhibits better performance in reconstructing the empirical network’s strength distribution and sequence and produces similar morphological patterns to the classic Maslov-Sneppen algorithm. Finally, we demonstrate how the choice of a network null model can influence weighted network inference through the example of weighted rich-club detection.

Network null models are so ubiquitous in network science that they have been integrated into classic measures such as the small-world coefficient [53, 76, 122] and the rich-club coefficient [28, 82]. In recent years, graph analysis of brain networks has moved away from earlier binarization procedures to instead consider the large range of edge weights provided by modern imaging and tracing technologies [10, 34, 49, 54, 77, 79, 118, 127]. Weighted graph measures are taking the place of their simpler binary counterparts, allowing researchers to probe the architectural properties characteristic of weighted brain networks [9, 76, 116] and the relationships between edge weights and their physical embedding [12, 88, 109]. Nevertheless, degree-preserving null models remain the most commonly used methods for network statistic normalization [53, 70, 78, 116, 118]. However, such models do not mitigate the possible effects of weighted degree (i.e. strength). Efforts towards statistical inference for next-generation connectomics should therefore focus on developing and evaluating strength-preserving network null models to adapt to the growing reliance on weighted networks in the field.

Several such null models that preserve a network’s strength sequence in addition to its degree sequence have already been developed. One class of such models is based on maximum-likelihood estimation of exponential random-graph models [41, 96, 104]. They allow unbiased sampling of networks satisfying degree and strength constraints on average, but not for each network realization. Furthermore, the generated networks do not preserve the empirical weight distribution [89]. Similarly, Roberts and colleagues designed a strength correction procedure that iteratively rescales the weights of a connectivity matrix to obtain any desired strength sequence [88]. An advantage of this method is that it can be easily combined with other constrained network randomization methods to generate surrogate networks embodying multiple constraints. For example, in [88] and [44], the procedure was used in combination with a geometry-preserving network null model. However, this procedure also does not maintain the empirical network’s weight distribution. Finally, in the case of directed networks, the Maslov-Sneppen algorithm has previously been adapted to also preserve the empirical strength distribution [82], but only of either the incoming *or* outgoing connections.

Here we consider a simple strength-preserving randomization procedure that builds upon a rewiring method already widely adopted in the field of network neuroscience [70]. This model is based on weight permutations constrained via simulated annealing [59, 60] and retains the empirical network’s size, density, degree and strength distribution and sequence [75]. We benchmarked this procedure against another strength-preserving randomization algorithm developed by Rubinov-Sporns [92] to characterize weighted, signed functional networks, and that was adapted to unsigned structural networks in the context of this study. We also compared its performance against the widely used Maslov-Sneppen degree-preserving rewiring. Across multiple performance metrics and data processing choices, we showed that the simulated annealing algorithm outperforms the two other null models in reconstructing the empirical network’s strength distribution and sequence. Furthermore, in many instances, the Rubinov-Sporns algorithm [92] exhibited a worse performance than the Maslov-Sneppen model, which does not explicitly account for strength. Therefore, in terms of strength preservation, the simulated annealing algorithm should be favoured to the other two models.

Moreover, by embedding null network realizations in a two-dimensional morphospace spanned by clustering and characteristic path length, we find that simulated annealing generates less variable morphospace representations than the Rubinov-Sporns algorithm across data processing choices. That is, null networks derived via simulated annealing are more stable in terms of the portion of the morphospace they occupy in relation to nulls generated via other methods, closely aligning with the Maslov-Sneppen null networks. Therefore, once again, the simulated annealing algorithm should probably be preferred to the other two surrogate models for the morphological fidelity of its randomization process and the proximity of the morphological patterns it generates with the well-studied random patterns of the Maslov-Sneppen algorithm.

More broadly, the present report showcases morphospaces as an informative yet under-utilized analytical step for understanding the evolution and final position of network architectures as they undergo randomization. Morphospaces have previously been used to relate topological patterns of complex networks in a common space and investigate potential dynamical design principles underlying their architectures [5, 6, 29, 44, 45]. Here, we introduce their use in the context of the assessment of null network ensemble variability and feature preservation. We chose clustering and characteristic path length as axes because of their common use in network neuroscience. However, any other global network statistic could have been considered. Nevertheless, this choice informs us on the effect it could have to use the simulated annealing algorithm as a null model for network inference when assessing the unexpectedness of a classic network measure: the small-world coefficient (i.e., ratio of normalized clustering to normalized characteristic path length) [53, 76, 122]. Across datasets and data processing choices, null networks generated via simulated annealing show similar clustering to Maslov-Sneppen nulls, but slightly shorter characteristic path length, making them slightly more stringent in assessing how unexpectedly low a network’s characteristic path length is, and therefore, more conservative in characterizing a network as exhibiting small-world architecture.

By considering null subsamples of increasing size, morphospaces also provide a way to quantify scaling behaviors. We find that global weighted network features rapidly converge with the number of generated nulls across all null models, indicating that only a relatively small number of nulls might be necessary for robust normalization of network statistics. Furthermore, using individual participant brain networks, we show that morphospace representation can constitute a tool for the assessment of identity preservation in surrogate networks: by comparing inter-individual morphospace distance in empirical and null networks, we find that simulated annealing provides the best preservation of participant identity.

We have shown how the simulated annealing algorithm can be applied to preserve degree and strength sequence, but in principle, other features and feature combinations could also be preserved. Using null models that embody a hierarchy of constraints is in fact a fundamental process in network neuroscience [12, 61, 88– 90, 119]. When considered in parallel, they allow us to distinguish the additional contribution of higher-order constraints from those inherited from more influential lower-order constraints. For example, geometric constraints can be readily implemented in the simulated annealing objective function, allowing us disentangle the effects of the network’s topology from those passively endowed by spatial embedding [89].

Similarly, other low-level influential network features such as clustering, modules, or motifs could be explored in constraint hierarchies using simulated annealing. Such attempts have been made in the past, in both various real-world networks and brain networks [74, 83, 89]. However, these models should be thoroughly tested for specific applications as certain feature combinations might involve optimization trade-offs. Furthermore, certain constraints, such as those imposed on clustering, have been reported to lead to practical breaking of the condition of ergodicity, i.e., architectures are formed during rewiring that cannot be destroyed in realistic time frames [26, 83]. Null models embodying more complex constraints might therefore benefit from other methods from statistical physics such as the Wang-Landau algorithm [26, 36, 121].

The present findings should be considered in the context of some methodological limitations. First, while there exists a broad range of edge weight quantification schemes in modern connectomics, we only focused on weight metrics based on streamline counts. However, we find that the performance of simulated annealing generalized to a broad range of real-world complex networks. Therefore, we hypothesize that it should also be the case for other connectome edge weight metrics. Second, while the datasets covered a broad range of possible processing choices, they did not allow to delineate their individual effects on null model performance. As tools for multiverse analysis in connectomics are developed to facilitate the isolation of specific processing choices across a range of pipelines [40], future work can increasingly interrogate how they affect connectomes and downstream network analysis, including null network generation.

In conclusion, we characterize and benchmark a simple simulated annealing method for generating strength sequence-preserving randomized null networks. We find 13 that this model outperforms other network randomization algorithms. Building on top of conventional rewiring procedures already common-place in network neuroscience, this algorithm allows for flexible constraint optimization. By simplifying the generation of hierarchies of preserved features, the simulated annealing procedure presented in this study has the potential to transform network inference, allowing greater insight into the principles that shape brain networks.

## METHODS

Code and data used to perform the analyses can be found at https://github.com/netneurolab/milisav_strength_nulls.

### Data acquisition and network reconstruction

All analyses were applied on two datasets acquired and preprocessed independently. Namely, we used structural connectivity measures derived from data collected at the Lausanne University Hospital [46] and as part of the Human Connectome Project (HCP) S900 release [117]. Together, these data span a variety of methodological choices, allowing us to asses the robustness of the results. These differences in methodology include the use of multiple parcellation resolutions in a structural and a functional atlas for the Lausanne and the HCP datasets, respectively.

#### Lausanne dataset

The Lausanne dataset consists of data collected in *n* = 70 healthy participants (age 28.8 ± 8.9 years old, 37% females) that were scanned at the Lausanne University Hospital in a 3-Tesla Siemens Trio MRI Scanner using a 32-channel head-coil [46]. The protocol included (1) a magnetization-prepared rapid acquisition gradient echo (MPRAGE) sequence sensitive to white/gray matter contrast (1 mm in-plane resolution, 1.2 mm slice thickness) and (2) a diffusion spectrum imaging (DSI) sequence (128 diffusion-weighted volumes and a single b0 volume, maximum b-value 8000 s/mm^2^, 2.2 *×* 2.2 *×* 3.0 mm voxel size). Informed written consent was provided by all participants in accordance with institutional guidelines and the protocol was approved by the Ethics Committee of Clinical Research of the Faculty of Biology and Medicine, University of Lausanne, Switzerland.

For each participant, structural connectivity networks were reconstructed using deterministic streamline tractography. Nodes were defined according to a multi-scale grey matter parcellation [21]. In the present work, we use a fine 1000 cortical regions parcellation and a relatively coarser 219 nodes resolution. The FreeSurfer version 5.0.0 open-source package was employed to segment white matter and grey matter from the MPRAGE volumes, whereas tools from the Connectome Mapper open-source software [31] were used to preprocess DSI data. For each white matter voxel, 32 streamline propagations were initiated per diffusion direction [123]. Structural connectivity was defined as the streamline density between node pairs, i.e., the number of streamlines between two regions normalized by the mean length of the streamlines and the mean surface area of the regions [47]. Additional information regarding MRI data preprocessing and network reconstruction is available at [46].

Group-representative structural networks were then generated to amplify signal-to-noise ratio using functions from the netneurotools open-source package (https://netneurotools.readthedocs.io/en/latest/index.html). We adopted a consensus approach that preserves (a) the mean density across participants and (b) the participant-level edge length distribution [13]. First, the cumulative edge length distribution across individual participants’ structural connectivity matrices is divided into *M* bins, with *M* corresponding to the average number of edges across participants. The edge occurring most frequently across participants is then selected within each bin, breaking ties by selecting the edge with the highest average weight. This procedure was performed separately for intra- and inter-hemispheric edges to ensure that the latter are not under-represented. The selected edges constitute the distance-dependent group-consensus structural network. Finally, the weight of each edge is computed as the mean across participants.

To examine the effect of network density on null model process time, we considered the 219 nodes resolution. We scaled the average number of intra- and inter-hemispheric edges across participants separately to generate a range of *M* values (i.e., 3602, 3962, 4356, 4794, 5276, 5706, 6162, 6550, 6952, 7346, 7824, 8206, 8596, 9006, 9410, 9862, 10324, 10742, 11180, 11564, 11978, 12460, 12874, 13310, 13622, 14108, 14492). These values were used in the distance-dependent consensus procedure and 100 null networks were generated per *M* value for each null model.

For the participant-level analyses, we fixed the density of all individual networks to that of the most sparse network by pruning connections with the lowest weights. This allows for unbiased inter-individual comparisons of global network statistics.

#### Human Connectome Project dataset

MRI data from n = 327 unrelated healthy participants (age 28.6 ± 3.73 years old, 55% females) were used to reconstruct structural connectivity networks and build a group-representative connectome [11, 97]. The participants were scanned at Washington University in the HCP’s custom 3-Tesla Siemens “Connectome Skyra” MRI scanner. The protocol included (1) a magnetization-14 prepared rapid acquisition gradient echo (MPRAGE) sequence (TR = 2400 ms, TE = 2.14 ms, FOV = 224 mm *×*224 mm, voxel size = 0.7 mm^3^, 256 slices) and (2) a spin-echo echo-planar imaging (EPI) sequence (TR = 5520 ms, TE = 89.5 ms, FOV = 210 mm *×*180 mm, voxel size = 1.25 mm^3^, b-value = 1000, 2000, and 3000 s/mm^2^, 270 diffusion directions, 18 b0 volumes). Informed written consent was provided by all participants and the protocol was approved by the Washington University Institutional Review Board. Additional information regarding data acquisition is available at [117].

The HCP minimal preprocessing pipelines [43] were applied to the MRI data and streamline tractography tools from the MRtrix3 open-source software [112] were used to reconstruct structural connectivity networks in individual participants from diffusion-weighted MRI (dMRI) data. The MPRAGE volume was segmented into white matter, grey matter, and cerebrospinal fluid to perform anatomically constrained tractography. Grey matter was divided according to the 400 and 800 cortical regions resolutions of the Schaefer functional parcellation [94]. The multi-shell multi-tissue constrained spherical deconvolution algorithm from MRtrix3 was used to generate fiber orientation distributions [25, 55]. The tractogram was initialized with 40 million streamlines and constrained with a maximum tract length of 250 and a fractional anisotropy cutoff of 0.06. A spherical deconvolution-informed filtering procedure (SIFT2) was then applied following Smith et al. [99] to estimate streamline-wise cross-section multipliers. Additional information regarding MRI data preprocessing and network reconstruction is available at [85].

A group-representative weighted structural connectivity matrix was then generated following the same consensus approach used for the Lausanne dataset. Finally, the weights of the group-consensus structural network were log-transformed and scaled between 0 and 1 to reduce variance in strength.

#### Drosophila

The drosophila connectome was reconstructed using 12 995 projection neuron images of the female Drosophila brain from the FlyCircuit 1.1 database [24, 98]. Imaged neurons were labeled with green fluorescent protein (GFP) using genetic mosaic analysis with a repressible cell marker (MARCM; [62]). GFP-labeled neurons were then delineated from whole brain three-dimensional images. Individual GFP-labeled neurons were coregistered to a female drosophila template brain using an affine transformation. Each neuronal soma was segmented and its nerve fiber skeleton was automatically traced with an algorithm using the soma’s center as the point of origin. The neurons were then divided into 49 local populations with distinct morphological and functional characteristics, which constituted the nodes of the network. These local populations consisted in 43 local processing units (LPUs) and 6 interconnecting units. An LPU was defined as a region with its own population of local interneurons whose fibers are circumscribed to this region. This analysis resulted in a weighted, directed network, with the weight of each edge defined as the number of neuron terminals between two populations. Additional information regarding the drosophila connectome reconstruction is available at [98].

#### Mouse

The mouse connectome was reconstructed using publicly available data from 461 tract-tracing experiments conducted in wild-type mice by the Allen Institute for Brain Science [79, 93]. Analyses were performed using a whole-brain 130 regions bilaterally symmetric parcellation designed from the Allen Mouse Brain Atlas [63, 79] and the Allen Developing Mouse Brain Atlas [111]. Each experiment consisted in an anterograde tracer injection into one of the 65 regions of the right hemisphere followed by whole-brain (intra-hemispheric and interhemispheric) imaging and mapping of axonal projections. Connection weights were defined as normalized connection densities (NCD), proportional to the number of axons projecting from one unit volume of the source region to one unit volume of the target region. Nine regions in each hemisphere were discarded due to the absence of experiments, resulting in a connectome composed of 112 nodes and spurious connections were removed using a probabilistic threshold (*P <* 0.01) based on expert visual assessment [79]. Hemispheric symmetry was assumed and the available interhemispheric projections were used to reconstruct the whole-brain connectome. As information on the tracer injections’ source and target sites was provided, this analysis resulted in a weighted, directed network. Additional information regarding the mouse connectome reconstruction is available at [79, 93].

#### Rat

The rat connectome was reconstructed using *>* 16 000 rat cortical association macroconnection (RCAM) histological reports from *>* 250 systematically curated references in the primary literature, publicly available in the Brain Architecture Knowledge Management System (BAMS; [15–18] Considered reports were based on experimental tract tracers transported anterogradely, retrogradely, or in both directions within axons. Analyses were performed using 73 cortical regions derived from the hierarchical Swanson-04 nomenclature [17, 110]. One of 8 ranked qualitative connection weights was assigned to each connection based on the reports with the most accurate and reliable tracer methodology, optimal injection sites, and highest anatomical density of tracer labeling (neuronal soma for source and axon ter-15 minals for target). Connection weights were then transformed to approximately logarithmically spaced weights in the range [0, 1]. As information on the tracer injections’ source and target sites was provided, this analysis resulted in a weighted, directed network. Additional information regarding the rat connectome reconstruction is available at [15].

#### Macaque

The macaque connectome was reconstructed using the macroconnectivity CoCoMac database [106], collating results from tract-tracing studies in macaque monkeys [95]. Analyses were performed using the combined Walker-von Bonin and Bailey (WBB47) atlas [105], containing 39 non-overlapping cortical regions. An anatomical tract was included between two brain regions if (1) it was reported in at least five studies in the database and (2) at least 66% of the reports were positive (i.e., detected its presence; [32]). Edges were weighted between 1 and 3 based on their average reported strength. As information on the tracer injections’ source and target sites was provided, this analysis resulted in a weighted, directed network. Additional information regarding the macaque connectome reconstruction is available at [95].

#### Complex weighted networks

To benchmark strength sequence-preserving null network models beyond the use-cases of connectomics, we use a diverse dataset of 37 real-world complex weighted networks. These networks were previously used by Oldham et al. [80] to compare centrality measure and build on top of another dataset of complex networks compiled by Ghasemian et al. [42]. All networks but one originate from the Index of Complex Networks (ICON; [27]). The final network is a human structural brain network derived from diffusion-weighted MRI data from the HCP (see [80] for more details). Overall, the network sizes range from 13 to 1707 nodes and their densities range from 0.3% to 78.31% of connections present. The dataset bridges a number of distinct data categories, with 28 networks representing social data, 1 network representing informational data, 2 networks representing biological data, 1 network representing economic data, and 5 networks representing transportation data.

### Null models

Two network null models for the strength sequence-preserving randomization of undirected weighted adjacency matrices were considered as part of the current report. Both algorithms also perfectly preserve the original edge weight distribution. The first model was developed by Rubinov & Sporns for signed functional brain networks [92] and adapted here to strictly positive structural networks. The second model implements a simulated annealing procedure. To our knowledge, it was introduced in [75] as a technique for strength sequence-preserving randomization and further explored in directed connectomes for the preservation of multiple network features in [89]. Python implementations of both algorithms have been made openly available as part of the netneurotools package (https://netneurotools.readthedocs.io/en/latest/api.html#module-netneurotools.networks).

The two strength sequence-preserving randomization models were benchmarked against the Maslov-Sneppen rewiring algorithm that randomly swaps pairs of edges, systematically disrupting the topology of the empirical network, while preserving network size (i.e., number of nodes), density (i.e., proportion of expressed edges), and degree sequence (i.e., number of edges incident to each node), but not strength sequence (sum of edge weights incident to each node) [70]. This model was implemented using the openly available *randmio_und_connected* function from the Python version of the Brain Connectivity Toolbox (https://github.com/aestrivex/bctpy) [91], specifying approximately 10 swaps per edge. This function has the additional advantage of maintaining connectedness in the rewired network, i.e., no node is disconnected from the rest of the network. This constraint is also met in both strength sequence-preserving null models, because they operate on networks rewired according to the Maslov-Sneppen algorithm, simply rearranging weights atop the randomized binary network scaffold to further maintain strength sequence.

The Rubinov-Sporns algorithm [92] consists of the following steps. First, the randomized network weights are instantiated with zeros *Âij* = 0. Second, the original edge weights are ranked by magnitude. Third, the edges of the randomized network are ranked by the expected weight magnitude *ê*_*ij*_ *∝* (*s*_*i*_ *−* Σ_*u*_ *Â* _*iu*_)(*s*_*j*_ *−* Σ_*u*_ *Âju*), where *s*_*i*_ is the original strength of node *i*. Fourth, a random edge is selected in the randomized network and its weight *Âij* is set to the original edge weight of the same rank. Finally, the associated edge and weight are removed from further consideration and the remaining original edge weights and randomized network edges are again ranked as described above. The procedure is repeated until every edge in the randomized network has been assigned one of the original weights.

The last null model employs simulated annealing to preserve the empirical network’s strength sequence. Simulated annealing is a stochastic search algorithm that approximates the global minimum of a given function [59, 60] using the Metropolis technique [72], controlled by the temperature parameter *T*. A high temperature regime allows the exploration of costly system configurations, whereas fine-tuned adjustments with smaller effects on the system cost are provided at lower temperatures. Initially, the simulated annealing algorithm is set 16 at a high temperature, preventing the process from getting stuck in local minima. Throughout the procedure, the system is slowly cooled down while descending along the optimization landscape, yielding increasingly limited uphill rearrangements. Here, we minimize the cost function *E* defined as the mean squared error (MSE) between the strength sequence vectors of the original and the randomized networks. When randomizing directed networks, we adapt the cost function by separately computing MSE for the in-strength and the out-strength and summing both results to obtain *E*. To optimize this function, random weight pairs are swapped. Reconfigurations were only accepted if they lowered the cost of the system or met the probabilistic Metropolis acceptance criterion: *r < exp*(*−*(*E′ −E*)*/T*), where *r ∼ U* (0, 1). The annealing schedule consisted in 100 stages of 10 000 permutations with an initial temperature of 1000, halved at each stage. Pseudocode detailing the complete strength sequence-preserving simulated annealing is available as Algorithm 1.

For the participant-level analysis, performance is evaluated at each stage and the annealing schedule is halted if performance reaches a predefined energy threshold of 0.0001 or runs for a maximum of 1000 stages. This threshold was chosen to ensure a uniform final energy distribution across nulls of different empirical networks and avoid inter-individual differences in downstream analyses due to differences in the quality of the null strength reconstruction.

### Performance metrics

- *MSE*. In the context of this study, mean squared error (MSE) is defined as:

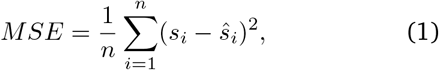

where *n* is the number of nodes in the network, *s*_*i*_ is the strength of node *i* and *ŝ*_*i*_ is the strength of node *i* in the randomized network.
- *Process time*. The process times reported in this study strictly correspond to CPU execution times of the randomization processes in seconds. They were computed as the differences between consecutive calls to the *process_time* function of the Python Standard Library time module. The calls to the *process_time* function were placed immediately before and after the calls to the randomization algorithms. To relate process time to the density of the network to be randomized, the process time of the Maslov-Sneppen rewiring was subtracted from the process time of the two strength-preserving randomization algorithms, because they both incorporate the procedure, which depends on the number of edges in the network.

### Topological features

- *Degree*. Degree corresponds to the number of edges incident on a node.
- *Strength*. Weighted degree, or strength, corresponds to the sum of edge weights incident on a node.
- *Clustering coefficient*. The weighted local clustering coefficient of a node corresponds to the mean “intensity” of triangles around a node. The clustering coefficient *C* of a node *u* can be defined as [81]:

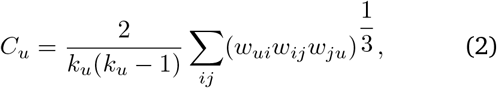

where *k*_*u*_ is the degree of node *u* and *w*_*ij*_ is the weight of the edge incident on nodes *i* and *j*, scaled by the largest weight in the network. It was computed using the openly available *clustering_coef_wu* function from the Python version of the Brain Connectivity Toolbox (https://github.com/aestrivex/bctpy) [91]. Prior to applying the function, the weights of the network were normalized to the range [0, 1] using the *weight_conversion* function from the Python version of the Brain Connectivity Toolbox (https://github.com/aestrivex/bctpy) [91]. The network average clustering coefficient was computed as the mean across weighted local clustering coefficients of all nodes in the network.
- *Characteristic path length*. The negative natural logarithm was used to map edge weights to lengths in all human connectomes, whereas an inverse transformation was used for all other networks to account for various weight ranges. Dijkstra’s algorithm [33] was then used to identify the sequence of unique edges spanning the minimum length between each node pair (i.e., shortest path). Shortest path lengths were then computed as the sums of traversed edges’ lengths. This procedure was implemented using the openly available *distance_wei* function from the Python version of the Brain Connectivity ToWWWolbox (https://github.com/aestrivex/bctpy) [91].

The characteristic path length of a network was finally computed as the average across all shortest path lengths in the network.

### Hub identification

Network hubs are a set of centrally embedded nodes [114]. When considering (weighted) degree as the centrality measure in question, the presence of hubs in a network is often detected based on the presence of a heavy-tail degree distribution. In contrast, an exponentially de-17 caying degree distribution is consistent with expectations from random networks [37].

Parametric modeling of degree distributions is often used to detect heavy-tailedness. However, these methods depend on subjective processing choices. Therefore, following Gajwani et al. [40], we opt for a non-parametric approach to facilitate comparison across datasets and null models [56].

Briefly, we assess the heavy-tailedness of a degree distribution by determining how heavy its right tail is in contrast to what is expected of the exponential distribution, i.e., whether it shows subexponential tail decay. To do so, we compute the empirical first and third quartiles, denoted as *Q*1 and *Q*3, respectively, along with the interquartile range (*IQR* = *Q*3 *− Q*1). The measure of “right-tailedness” is then defined as the probability that a randomly selected observation from the distribution exceeds the value represented by *Q*3 + 3 *∗ IQR* (i.e., *p*_*R*_(*X*) = *P* (*X > Q*3+3*∗IQR*)), a widely used definition of outliers [71]. This computed probability can be contrasted with the right-tailedness of the exponential distribution (*X ∼ e*^*−λx*^), which remains constant across different shape parameters (*λ*). Specifically, *p*_*R*_(*X*) *≈* 0.009 [56] and heavy-tailedness is detected if the empirical *p*_*R*_ exceeds this analytic threshold.

Hubs are defined as the outliers (values exceeding *Q*3+3*∗IQR*) and the z-scored Rand index is used [87] to measure the similarity of the partitions of network nodes into hubs and non-hubs between empirical networks and their randomized nulls.

### Weighted rich-club detection

A rich-club is defined as a a set of high-degree nodes (rich nodes) that share more connections than expected by chance [28, 129]. In this study, to further account for the relative strength of rich links, i.e., links connecting rich nodes, we consider a weighted measure of the rich-club phenomenon, the weighted rich-club coefficient [82]:

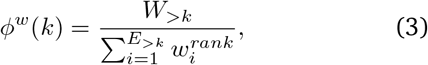

where *W*_*>k*_ is the total weight of the rich links, *E*_*>k*_ is the number of rich links, and 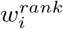 is the weight of the network edge at rank *i*, given a ranking by weight. Rich nodes are defined as nodes with degree *>*= *k* and a range of *k* values are considered.

In other words, the weighted rich-club coefficient measures the proportion of weight that rich nodes share in relation to the overall amount they could potentially share if linked by the network’s most robust connections.

Considering that high-degree nodes have a higher chance of being interconnected, the (weighted) rich-club coefficient is typically normalized against a null rich-club coefficient computed in a population of randomized null networks [82, 116]. While the classic Maslov-Sneppen degree-preserving rewiring [70] is often used to build the null network population even when computing the weighted rich-club coefficient, it does not account for the effect of weighted degree sequence on the weighted rich-club phenomena.

To contrast inferences obtained using different null models, we compute the weighted rich-club coefficient in each of the 10 000 null networks generated for each null model under study. We then compute the normalized weighted rich-club coefficient as:

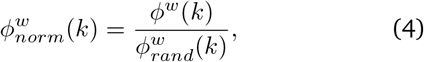

with 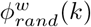 the mean weighted rich-club coefficient across the null network population.

Finally, to assess the statistical significance of the weighted rich-club phenomenon, a p-value is derived as the proportion of null *ϕ*^*w*^(*k*) that are greater than the empirical *ϕ*^*w*^(*k*).

## SUPPLEMENTARY INFORMATION

### Null model calibration

Scatter plots of strengths in the empirical and the randomized networks provide additional information about individual null model calibration and bias, i.e., the fidelity of its behavior across the range of data under consideration. For example, in the HCP dataset (Fig. 2a, S4), the Rubinov-Sporns algorithm appears to systematically underestimate low strengths and overestimating high strengths, as shown by the preponderance of data points below the identity line for lower empirical strength values and above the identity line for higher empirical strength values. By comparison, the Maslov-Sneppen algorithm shows a more even spread of points around the identity line, and even slightly outperforms the Rubinov-Sporns algorithm in terms of mean squared error (MSE; Fig. S1) in the low resolution HCP network (*p <* 10^*−*22^, effect size = 54.08%, two-tailed, Wilcoxon–Mann–Whitney two-sample rank-sum test). Importantly, the simulated annealing algorithm yields data points which perfectly align to the identity line, indicative of an unbiased fit. In contrast, in the Lausanne dataset (Fig. S4), the Maslov-Sneppen model underestimates high strengths, with most data points falling under the identity line at high empirical strength values. Conversely, the Rubinov-Sporns algorithm and the simulated annealing algorithm exhibit a number of data points above the identity line. In the case of simulated annealing, a small number of low strengths are over-estimated. Importantly, these errors do not appear to be systematic, as they affect different regions in different null realizations. Therefore, screening strength preservation results for each null realization could avoid introducing a bias towards the overestimation of low strengths in the null ensemble without incurring a considerably longer processing time. Furthermore, these low-strength outliers only have a small effect on simulated annealing performance, as shown by the high correlation coefficients obtained.

### Alternative objective function

To mitigate the small reconstruction errors observed in the Lausanne dataset using simulated annealing, we consider a different objective function that penalizes large individual errors: maximum absolute error. We explore the effect of this objective function in 100 null networks for each empirical network. We find that low-strength outliers are less apparent but that this effect is achieved at the detriment of the overall fit of the model (Fig. S2). For all networks, the strength sequence preservation is considerably worse as assessed using Spearman rank-order correlation (LAU - low res: *p <* 10^*−*33^, effect size = 85.10%, LAU - high res: *p <* 10^*−*65^, effect size = 100%, HCP - low res: *p <* 10^*−*68^, effect size = 100%, HCP - high res: *p <* 10^*−*66^, effect size = 100%, two-tailed, Wilcoxon–Mann–Whitney two-sample rank-sum test).

### Log-transformation

Our assessment of the quality of strength reconstruction revealed a slight tendency of the simulated annealing algorithm to overestimate low strengths in networks with a heavily right-skewed weight distribution. This could be due to low-strength nodes having lower degrees and therefore providing fewer “degrees of freedom” in matching their original strength through weight permutations. We posit that the small discrepancies in strength reconstruction observed between datasets might be due to a simple but potentially influential processing step: log-transformation, which was applied to the HCP dataset but not the Lausanne dataset. This common practice consists in taking the logarithm of the network’s edge weights and scaling them between 0 and 1 [11, 85]. Weight distributions of physical connections between neural elements have often been described as approximately log-normal across scales and species [7, 20, 69]. Log-transformation is therefore believed to bring a network’s weight distribution closer to normality, making it more amenable to downstream analyses. In the case of the simulated annealing algorithm, correcting the strong skewness of connectome weight distributions might allow for more degrees of freedom in reconstructing the strength sequence, consequently leading to faster convergence. In line with this hypothesis, we find that applying the log-transformation to the Lausanne dataset leads to better solutions (LAU low res: *p <* 10^*−*26^, effect size = 93.31%, LAU high res: *p <* 10^*−*33^, effect size = 100%) of similar quality to those obtained with the HCP dataset (Fig. S3).

### Morphospace trajectories

A question that might arise when using iterative optimization methods such as simulated annealing is how dependent the solution is on the number of iterations. Once again, morphospace representation can be used to address this question. Namely, it allows us to ask how much do global architectural features of null networks change as a function of optimization performance or number of steps taken. As an example, we consider the low-resolution empirical networks of the Lausanne and HCP dataset and we track the trajectory of a null network through morphospace during optimization (Fig. S10a,d). We find that while early optimization steps result in big leaps in the morphospace, i.e. highly variable solutions depending on the moment the process is stopped, displacements quickly localize to a constrained portion of the morphospace. Correspondingly, we observe a rapid transition in performance relatively early in the simulated annealing process (Fig. S10c,f). This indicates that simulated annealing reaches a high level of performance early on and that the following minor increases in performance do not result in important structural changes for the null networks. In other words, each null network, based on the simulated annealing procedure’s random initialization, has a predetermined place in the morphospace. This is further confirmed by aggregating morphospace positions and performance (in terms of MSE) across 100 null trajectories of 100 annealing stages (Fig. S10b,e). Indeed, we find a clear dichotomy between an area of low-energy and an area of high-energy. Therefore, tracking performance and variability in the null network features under consideration for a subset of nulls could be very useful in establishing an iteration threshold or performance target to ultimately reduce the duration of the null networks’ generation, while still preserving a good approximation.

### Weighted rich-club inference and geometry

In section “The weighted rich-club phenomenon”, we have previously shown that using simulated annealing-derived null networks yields larger normalized rich-club ratios than using Rubinov-Sporns or Maslov-Sneppen randomization (Fig. 4, left, S11, top; *p <* 0.01 for all two-tailed, Wilcoxon–Mann–Whitney two-sample ranksum tests). We posited that the differences in normalized rich-club coefficient observed between models was due to an overestimation of strength in high-degree nodes. Here, we verify this hypothesis and further relate this phenomenon to the prevalence of long-distance connections between hubs [115] and the exponential decay of connection weights with physical distance in structural brain networks [88, 109]. First, we consider the difference between the normalized rich-club coefficient obtained using simulated annealing and that obtained using Maslov-Sneppen rewiring. We relate this measure to median average weight (ratio of strength to degree) across the rich nodes. We find a strong negative relationship between the two measures (LAU - low res: *ρ* = *−*0.96, *p <* 10^*−*27^, LAU high res: *ρ* = *−*0.70, *p <* 10^*−*11^, HCP low res: *ρ* = *−*0.99, *p <* 10^*−*47^, HCP high res: *ρ* = *−*0.96, *p <* 10^*−*41^). This indicates that the difference in normalized rich-club coefficient between the two models might indeed be due to an overestimation of strength in high-degree nodes by the Maslov-Sneppen rewiring (Fig. S11; middle). We then relate median average weight to the median Euclidean distance across rich connections. Again, we find strong negative relationships between the two (LAU - low res: *ρ* = *−*0.91, *p <* 10^*−*18^, LAU high res: *ρ* = *−*0.93, *p <* 10^*−*32^, HCP low res: *ρ* = *−*0.87, *p <* 10^*−*19^, HCP high res: *ρ* = *−*0.62, *p <* 10^*−*8^), indicative of the previous effect possibly being due to a preponderance of low-weight long-range connections between rich nodes (S11; bottom).

**Figure S1.**
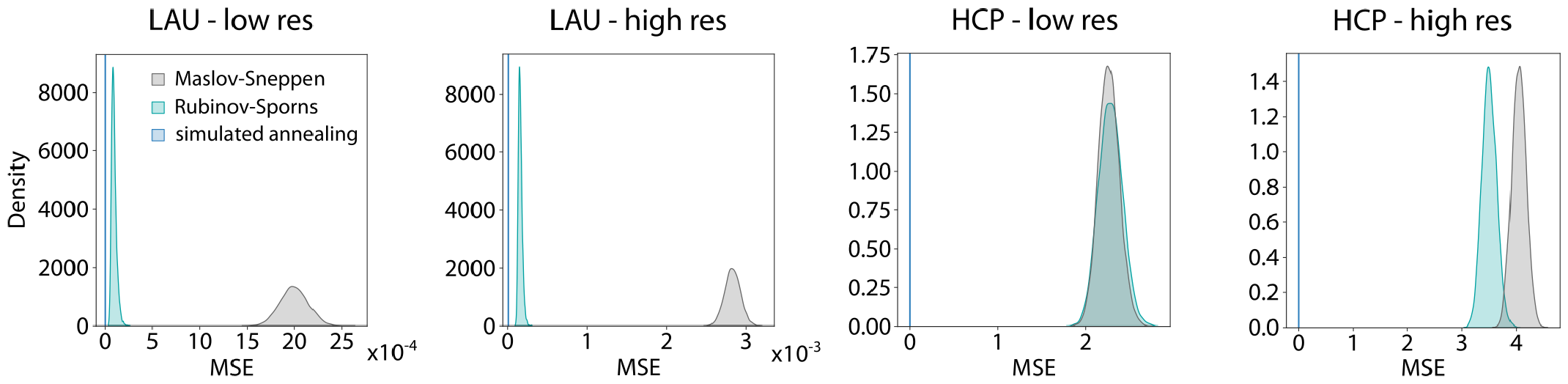
Benchmarking strength sequence preservation MSE. Density plots representing mean squared error between the strength sequences of the empirical and the randomized networks for the Maslov-Sneppen algorithm (grey), the Rubinov-Sporns algorithm (teal) and the simulated annealing algorithm (blue). Due to low variability, we only show the mean MSE across null realizations for the simulated annealing algorithm.

**Figure S2.**
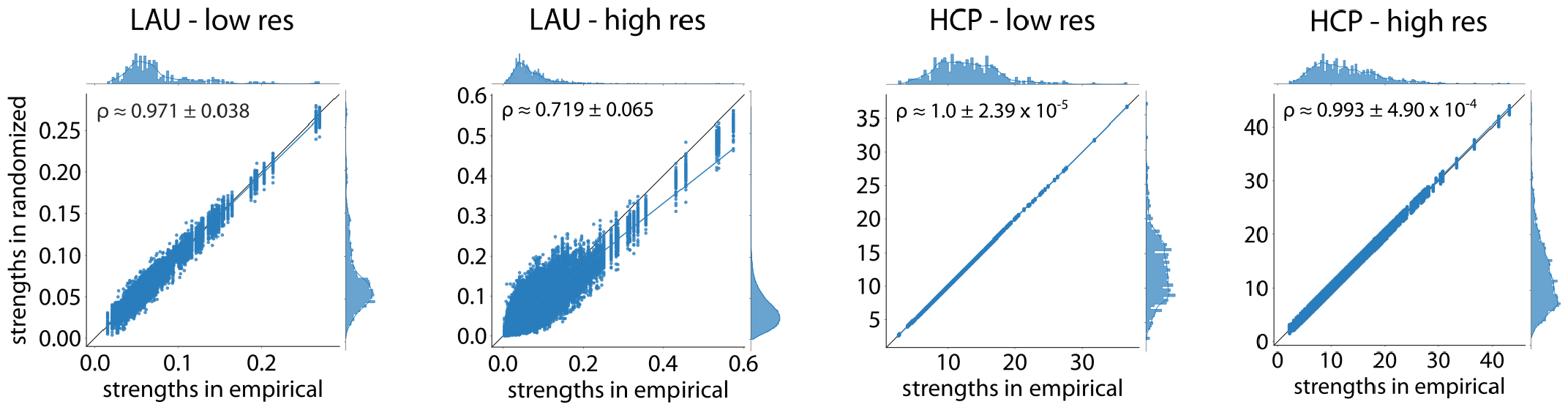
Benchmarking strength sequence preservation maximum absolute error. Scatter plots of strengths of the empirical (abscissa) and 100 null networks (ordinate) obtained using simulated annealing with the maximum absolute error objective function. Each point represents a brain region. Marginal distribution histograms are shown on the top and right axes. Mean and standard deviation across 100 Spearman rank-order correlation coefficients are provided as insets. Linear regression lines (blue) were added for visualization purposes. The identity line (black) is provided as reference.

**Figure S3.**
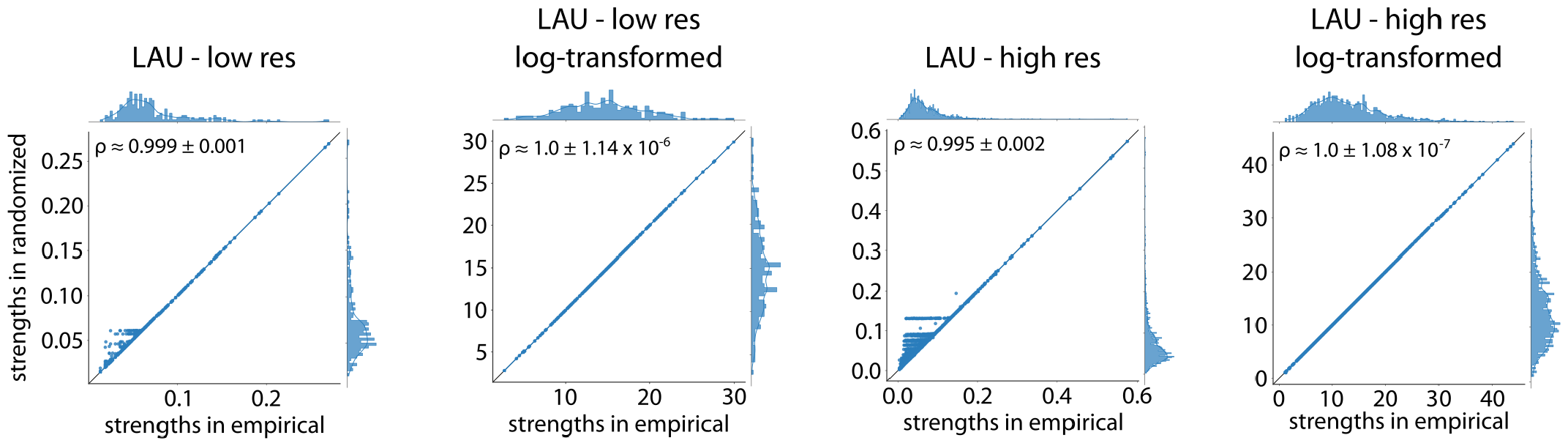
Benchmarking strength sequence preservation log-transformation. Scatter plots of strengths of the empirical (abscissa) and 100 null networks (ordinate) for the empirical and the log-transformed Lausanne networks. Each point represents a brain region. Marginal distribution histograms are shown on the top and right axes. Mean and standard deviation across 100 Spearman rank-order correlation coefficients are provided as insets. Linear regression lines (blue) were added for visualization purposes. The identity line (black) is provided as reference.

**Figure S4.**
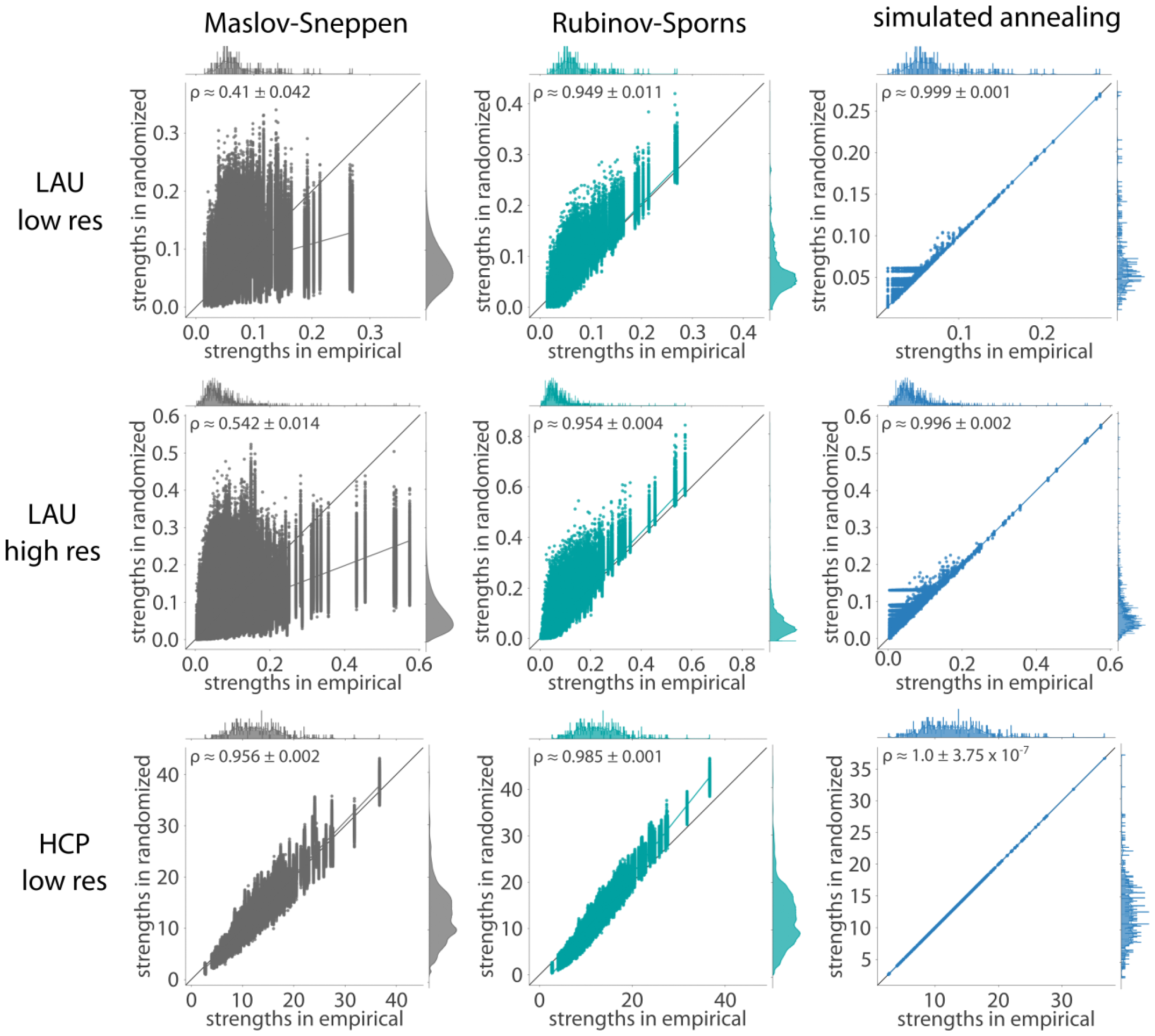
Benchmarking strength sequence preservation. Scatter plots of strengths of the empirical (abscissa) and randomized (ordinate) networks for all 10 000 null networks, where each point represents a brain region. Marginal distribution histograms are shown on the top and right axes. Mean and standard deviation across 10 000 Spearman rank-order correlation coefficients are provided as insets. Data points and histograms appear in grey for the Maslov-Sneppen algorithm, teal for the Rubinov-Sporns algorithm, and blue for the simulated annealing algorithm. Linear regression lines (colored) were added for visualization purposes. The identity line (black) is provided as reference.

**Figure S5.**
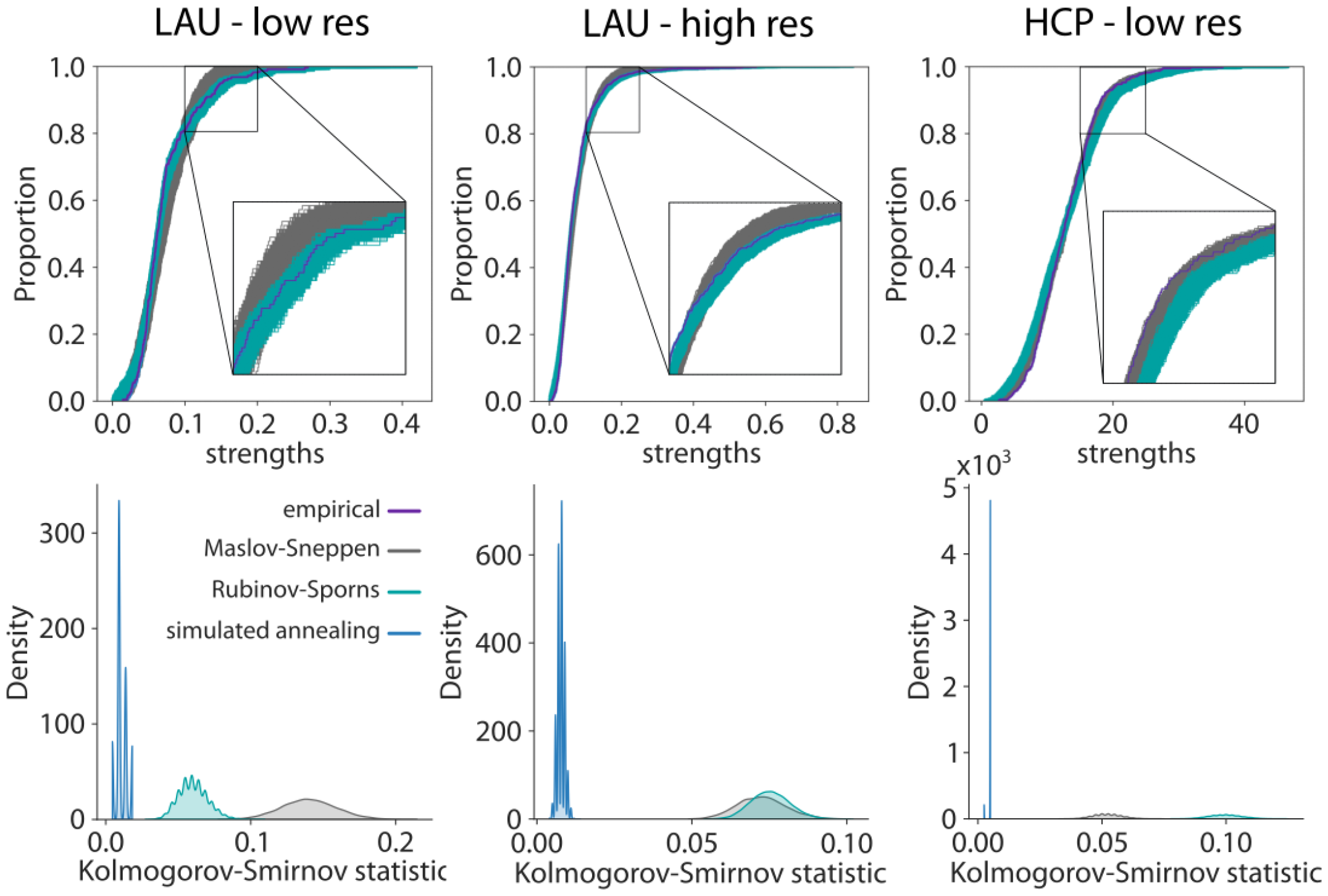
Benchmarking strength distribution preservation. Strength cumulative distribution functions (top) and density plots representing Kolmogorov-Smirnov statistics obtained by comparing the strength distribution of the empirical network with that of the randomized networks (bottom). Cumulative and probability density function curves are shown in grey for the Maslov-Sneppen algorithm, teal for the Rubinov-Sporns algorithm, and blue for the simulated annealing algorithm. The original cumulative distribution function is depicted in indigo and almost perfectly overlays all 10 000 cumulative distribution functions obtained via simulated annealing, effectively hiding them.

**Figure S6.**
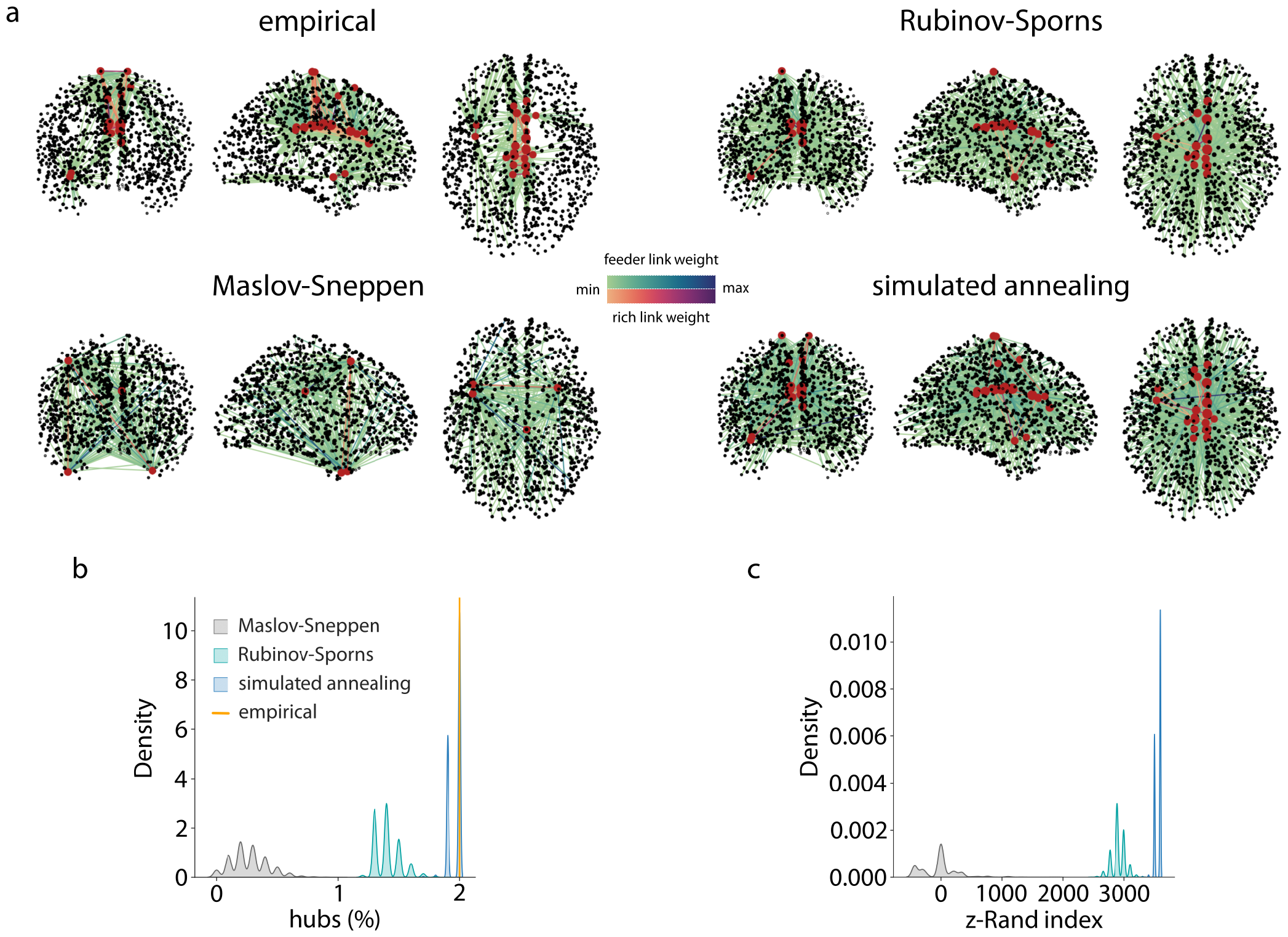
Hubs. (a) Brain plots representing hubs (red points) identified in the high resolution Lausanne empirical network (top left) and example Maslov-Sneppen (bottom left), Rubinov-Sporns (top right) and simulated annealing (bottom right) null networks. Feeder links (connections between hubs and non-hubs) are colored by weight based on the green-blue colormap, whereas rich links (connections between hubs) are colored by weight based on the red-indigo colormap. Displaying rich/feeder links shows that the network is rewired by all the algorithms, but the hubs are only preserved by the strength-preserving null models (Rubinov-Sporns and simulated annealing). (b) Density plots representing percentage of hubs in the empirical (yellow line) and across null networks (Maslov-Sneppen in grey, Rubinov-Sporns in teal, and simulated annealing in blue). (c) Density plots representing z-scored Rand indices between the empirical and the null hub assignments.

**Figure S7.**
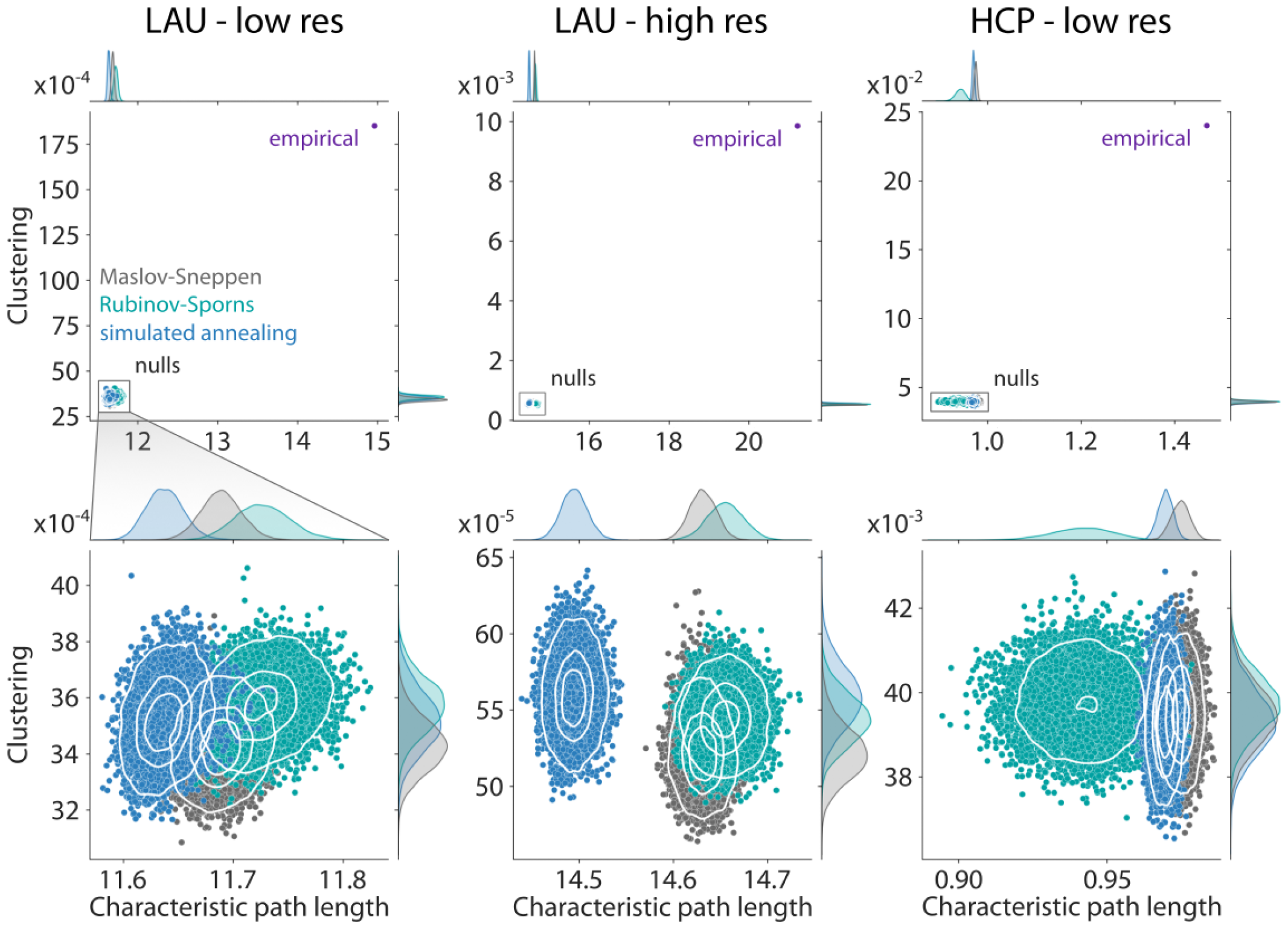
Morphospaces of null network ensembles. (a) Morphospaces spanned by characteristic path length and clustering. Marginal distribution histograms are shown on the top and right axes. Data points corresponding to randomized null networks generated by the simulated annealing algorithm appear in blue; those resulting from the Rubinov-Sporns algorithm appear in teal; and Maslov-Sneppen rewired networks are shown in grey. The empirical group-consensus structural network is depicted in indigo. The bottom panel consists in a zoomed-in view of the clusters of randomized networks appearing in the top panel. Contour levels are drawn using a Gaussian kernel density estimate and delineate iso-proportions of the density.

**Figure S8.**
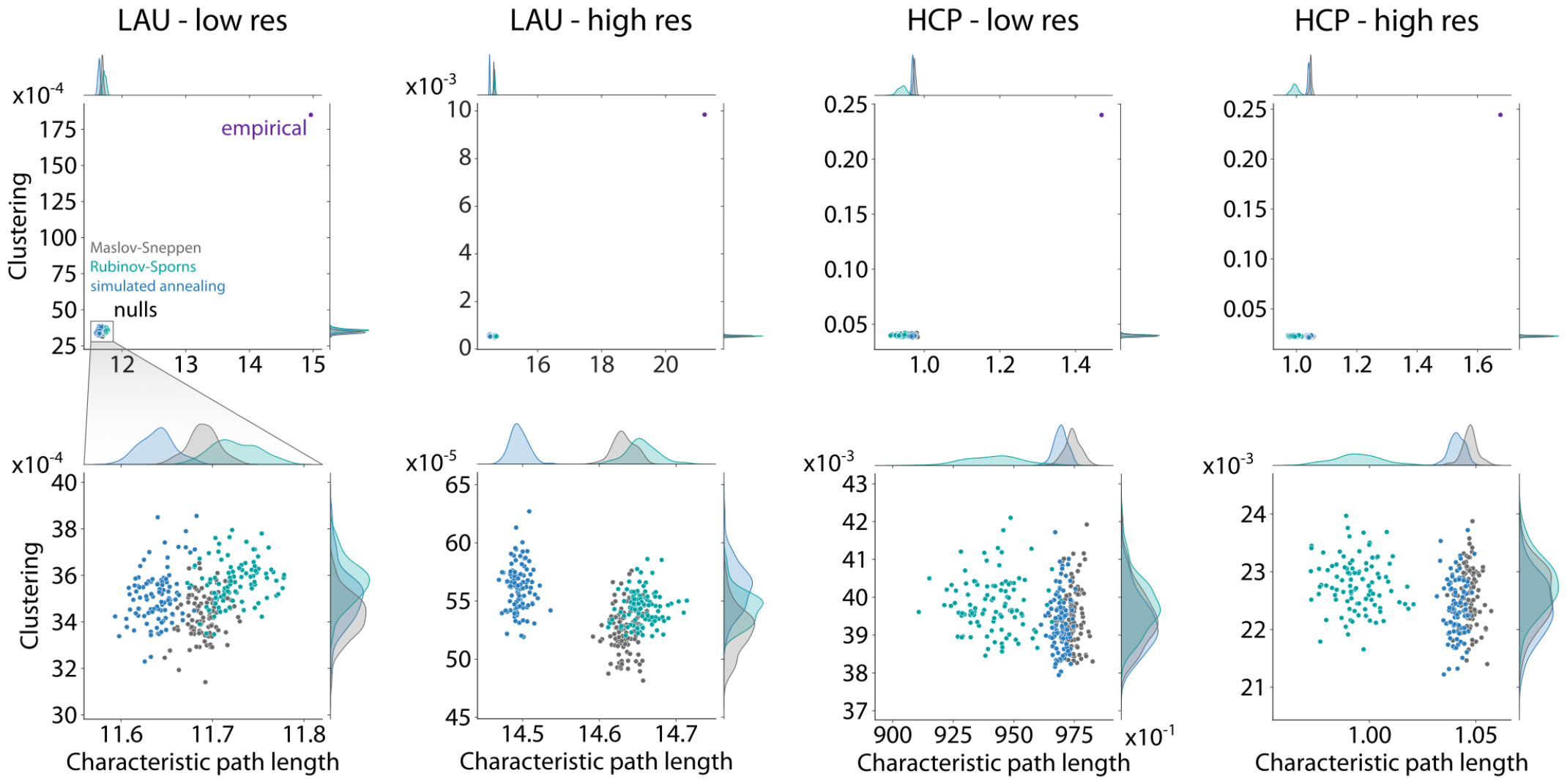
Morphospaces - 100 nulls. Morphospaces spanned by characteristic path length and clustering for a subset of 100 nulls. Marginal distribution histograms are shown on the top and right axes. Data points corresponding to randomized null networks generated by the simulated annealing algorithm appear in blue; those resulting from the Rubinov-Sporns algorithm appear in teal; and Maslov-Sneppen rewired networks are shown in grey. The empirical group-consensus structural networks are depicted in indigo. The bottom row consists in a zoomed-in view of the clusters of randomized networks appearing in the top row.

**Figure S9.**
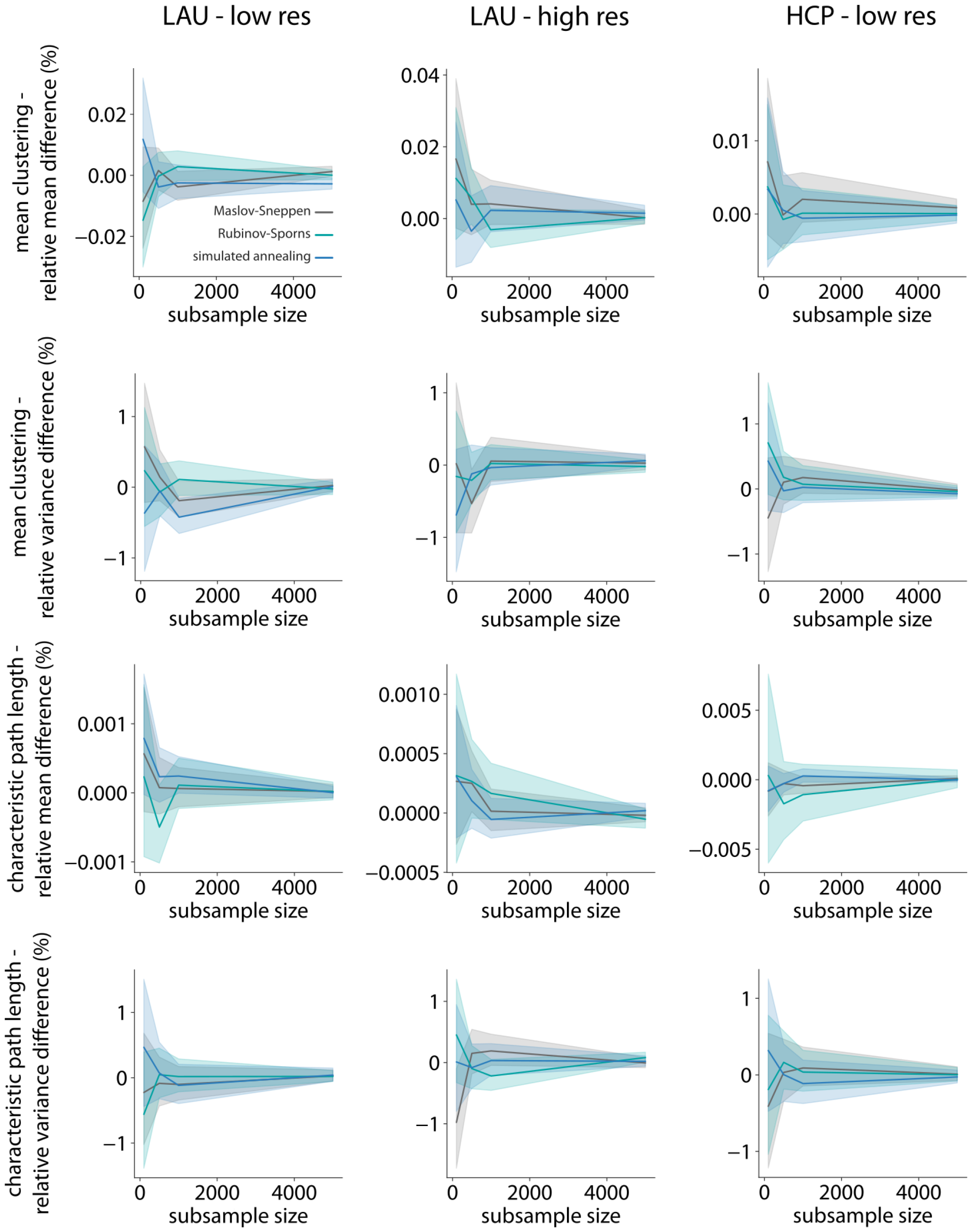
Morphospace scaling behavior. Trajectories of relative difference in mean clustering (first row), clustering variance (second row), mean characteristic path length (third row) and characteristic path length variance (fourth row) between the full null population (*N* = 10 000) and subsamples of increasing size (*n* ∈ {100, 500, 1000, 5000}). Colored lines and shaded bands represent mean and 95% bootstrapped confidence interval (1000 samples).

**Figure S10.**
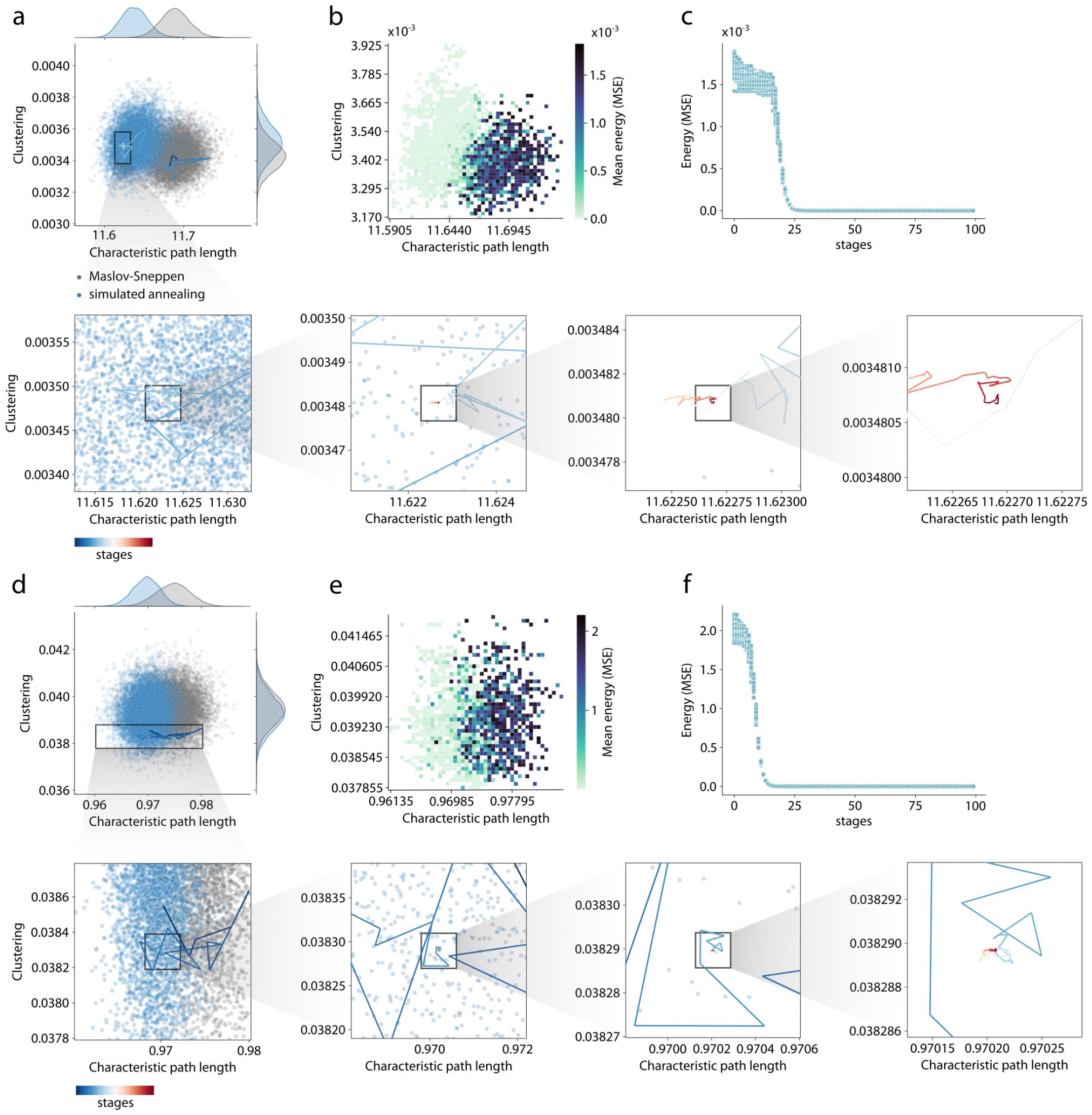
Morphospace trajectories. (a,d) Example annealing trajectory through a morphospace spanned by characteristic path length and clustering for the low-resolution Lausanne (a) and HCP (d) networks. Marginal distribution histograms are shown on the top and right axes. Data points corresponding to randomized null networks generated by the simulated annealing algorithm appear in blue and Maslov-Sneppen rewired networks are shown in grey. The trajectory is represented by a line colored from blue (early stages) to red (late stages). (b,e) Morphospace colored by mean energy aggregating 100 null network stages per annealing schedule across 100 nulls. (c,f) Optimization trajectories (energy as a function of annealing stage) for 100 null networks.

**Figure S11.**
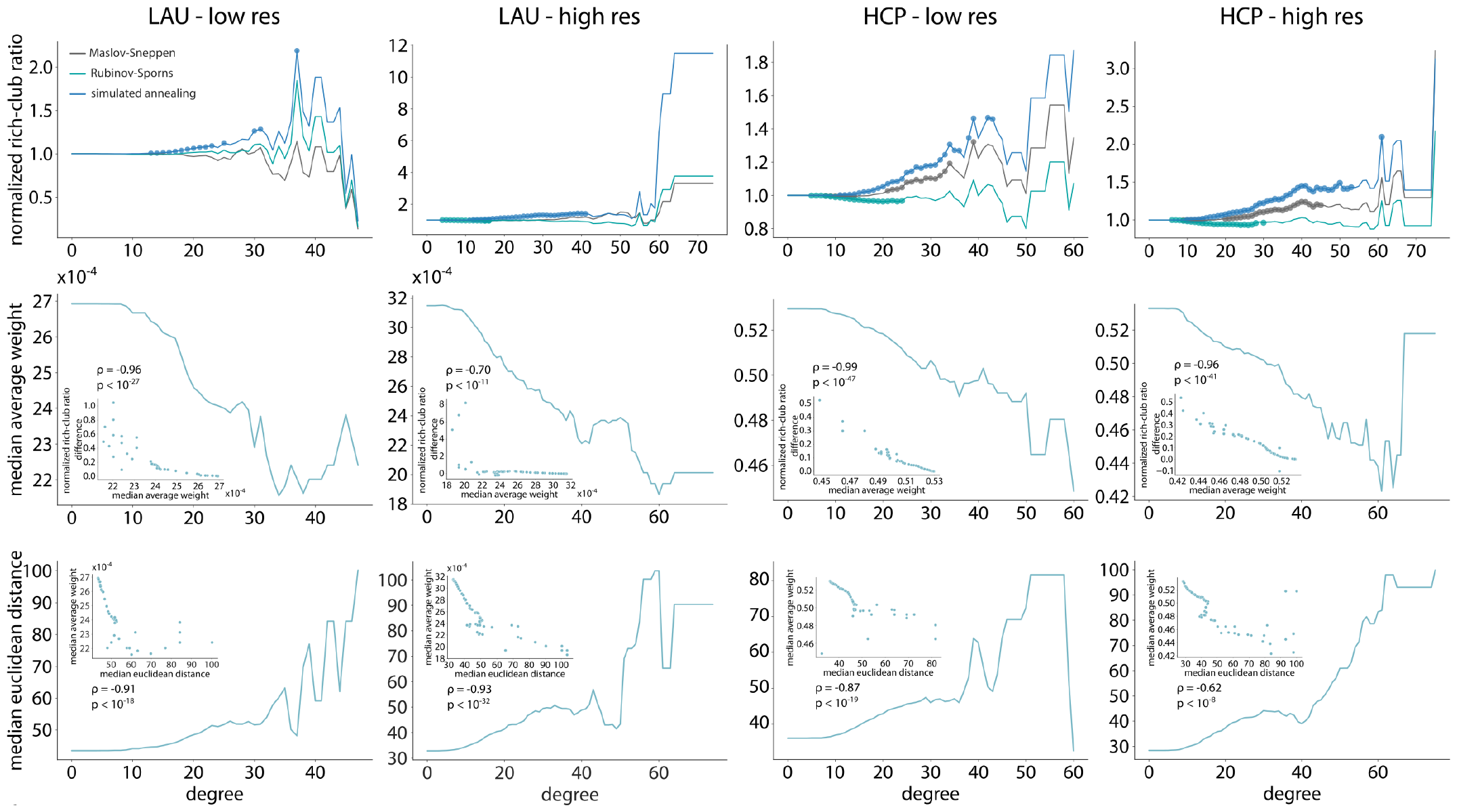
The weighted rich-club phenomenon. Top: Normalized rich-club ratio as a function of the degree threshold used to define rich nodes. Lines are colored by the null algorithm used (Maslov-Sneppen in grey, Rubinov-Sporns in teal, and simulated annealing in blue). Colored points indicate significance at the Bonferroni-corrected threshold of *p <* 0.05. Middle: Median average weight of the rich nodes as a function of the degree threshold used to define them. Inset: Relationship between the normalized rich-club ratio difference (simulated annealing-derived Maslov-Sneppen-derived) and the median average weight of the rich nodes. Spearman correlation coefficient and the resulting p-value are indicated above. Bottom: Median Euclidean distance of rich links as a function of the degree threshold used to define them. Inset: Relationship between the normalized rich-club ratio difference (simulated annealing-derived - Maslov-Sneppen-derived) and the median Euclidean distance of rich links. Spearman correlation coefficient and the resulting p-value are indicated below.

**Figure S12.**
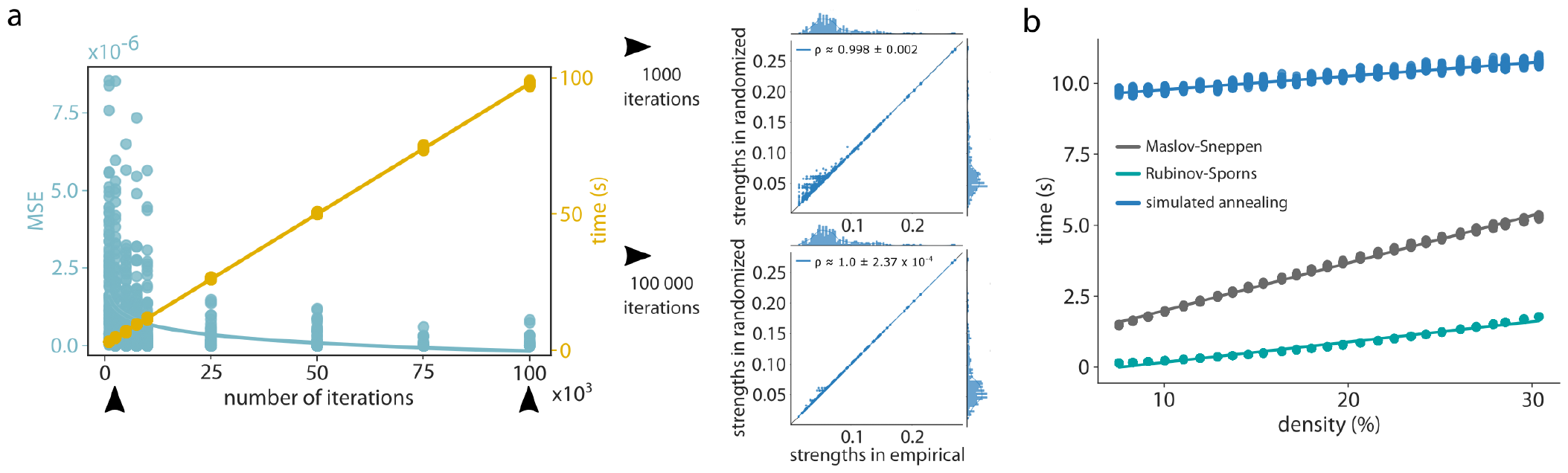
Computational cost-performance tradeoff of the simulated annealing procedure. All analyses were carried out on the low resolution group-representative connectome of the Lausanne dataset. (a) Left: MSE (*e*) decays logarithmically as a function of the number of iterations per annealing stage (*n*) while process time (*t*) increases linearly. The blue line corresponds to the least squares-fitted function *e* = *−*3.82 *×* 10^*−*7^*log*(*n*) + 4.22 *×* 10^*−*6^, *R*^2^ = 0.22. The yellow line corresponds to the least squares-fitted function *t* = 9.6 *×* 10^*−*4^*n* + 2.12, *R*^2^ *≈* 1.0 Right: Scatter plots of strengths of the empirical (abscissa) and randomized (ordinate) networks for all 100 null networks generated via annealing stages implementing 1000 (top) and 100 000 iterations (bottom). Mean and standard deviation across 100 Spearman rank-order correlation coefficients are provided as insets. Linear regression lines (blue) were added for visualization purposes. The identity line (black) is provided as reference. Interestingly, when the algorithm is allowed to run for more iterations, even the minor inaccuracies observed in the Lausanne dataset for low strength nodes (previously shown in Fig. S4) are completely “ironed out”, with a near-perfect reconstruction of the empirical strength sequence. (b) Linear relationships between process time (*t*) and density (*d*) for all three randomization algorithms. The blue line (simulated annealing) corresponds to the least squares-fitted function *t* = 0.0477*d* +9.3, *R*^2^ = 0.95. The teal line (Rubinov-Sporns) corresponds to the least squares-fitted function *t* = 0.0715*d −* 0.54, *R*^2^ = 0.98. The grey line (Maslov-Sneppen) corresponds to the least squares-fitted function *t* = 0.1674*d* + 0.32, *R*^2^ *≈* 1.0. Note that the duration of the Maslov-Sneppen rewiring was subtracted from that of the Rubinov-Sporns and simulated annealing randomizations.

**Figure S13.**
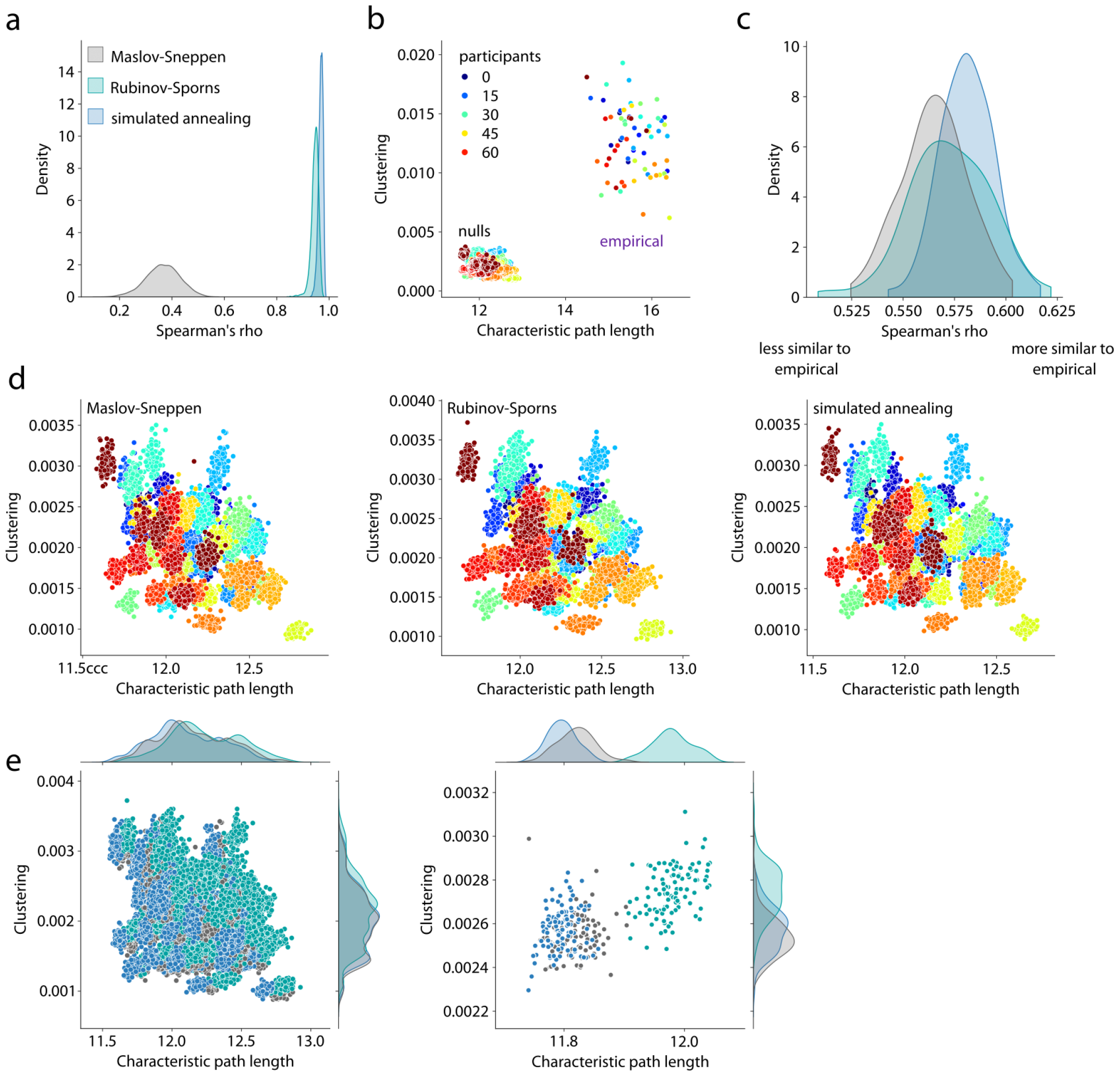
Strength-preserving randomization in individual networks. (a) Density plot representing Spearman correlation coefficients between strength sequences in empirical and randomized networks derived using the Maslov-Sneppen algorithm (grey), Rubinov-Sporns algorithm (teal), and simulated annealing algorithm (blue). (b) Morphospace spanned by characteristic path length and clustering in which 69 empirical connectomes and a total of 300 null networks per connectome were embedded. Networks are colored by participant. (c) Density plot representing Spearman correlation coefficients between empirical and null sets of Euclidean distances between participants across 100 nulls for each randomization algorithm. (d) Algorithm-wise morphospaces of null networks colored by participant. (e) Left: All null networks embedded in the morphospace. Right: Example null network morphospace for a single participant. Marginal distribution histograms are shown on the top and right axes. Data points are colored by the randomization algorithm used to generate them.

